# EMDiffuse: a diffusion-based deep learning method augmenting ultrastructural imaging and volume electron microscopy

**DOI:** 10.1101/2023.07.12.548636

**Authors:** Chixiang Lu, Kai Chen, Heng Qiu, Xiaojun Chen, Gu Chen, Xiaojuan Qi, Haibo Jiang

**Author notes:** Correspondence: Haibo Jiang, and Xiaojuan Qi.

## Abstract

Electron microscopy (EM) revolutionized the way to visualize cellular ultrastructure. Volume EM (vEM) has further broadened its three-dimensional nanoscale imaging capacity. However, intrinsic trade-offs between imaging speed and quality of EM restrict the attainable imaging area and volume. Isotropic imaging with vEM for large biological volumes remains unachievable. Here we developed EMDiffuse, a suite of algorithms designed to enhance EM and vEM capabilities, leveraging the cutting-edge image generation diffusion model. EMDiffuse demonstrates outstanding denoising and super-resolution performance, generates realistic predictions without unwarranted smoothness, improves predictions’ resolution by ∼30%, and exhibits excellent transferability by taking only one pair of images to fine-tune. EMDiffuse also pioneers the isotropic vEM reconstruction task, generating isotropic volume similar to that obtained using advanced FIB-SEM even in the absence of isotropic training data. We demonstrated the robustness of EMDiffuse by generating isotropic volumes from six public datasets obtained from different vEM techniques and instruments. The generated isotropic volume enables accurate organelle reconstruction, making 3D nanoscale ultrastructure analysis faster and more accessible and extending such capability to larger volumes. More importantly, EMDiffuse features self-assessment functionalities and guarantees reliable predictions for all tasks. We envision EMDiffuse to pave the way for more in-depth investigations into the intricate subcellular nanoscale structures within large areas and volumes of biological systems.

## Main

Electron microscopy (EM) is an essential tool for obtaining high-resolution images of biological specimens and has made a tremendous impact on cell biology, revealing highly complex cellular structures at the nanometer scale. Over the decades, EM has catalyzed numerous breakthroughs in life sciences, such as discovering novel cellular organelles, elucidating membrane structures, and generating detailed visualization of the macromolecular complexes^1-3^. Volume electron microscopy (vEM), a more recent innovation, arose from overcoming limitations of conventional two-dimensional electron microscopy techniques in studying 3D cellular structures^4-7^. vEM techniques such as serial section-based tomography techniques with transmission electron microscopy (TEM) and scanning electron microscopy (SEM), serial block-face SEM (SBF-SEM), and focused ion beam SEM (FIB-SEM) have facilitated investigations of 3D structures in cells, tissues and even small model organisms at nanometer resolution. Particularly, recent efforts in mapping brain connectomics with serial section-based vEM techniques^8-12^ and the developments of enhanced FIB-SEM for the acquisition of isotropic vEM data with improved robustness and throughput^13-15^ have further reinvigorated the field of EM and its applications.

However, the potential of EM and vEM is hindered by intrinsic limitations. It is essential to achieve a sufficient signal-to-noise ratio (SNR) to accurately map nanoscale cellular structures, which will inevitably reduce the imaging speed and prolong the dwell time per pixel, restricting the area and volume of imaging. Robust isotropic vEM also requires advanced instrumentation, such as enhanced FIB-SEM, which is only available at certain institutions or specialized centers. More importantly, the dogma of vEM is no viable method for generating isotropic datasets from large volumes. Currently, vEM is either limited to producing isotropic data from volumes of up to approximately 300 μm × 300 μm × 300 μm^16^, using state-of-the-art enhanced FIB-SEM, or capturing larger volumes aiming for cubic millimeter scale through serial section-based techniques^10, 17^, which, however, suffers from unsatisfactory axial resolution due to the limited section thickness of diamond knife cuts.

In recent years, computational methods and deep learning techniques have been applied to address the technical limitations of microscopy methods to expedite imaging processes and enhance the image quality of both light microscopy^18-21^ and electron microscopy^22-24^. Significant advances have been achieved in denoising and super-resolution tasks with deep learning techniques^23, 25-28^ However, three major limitations still exist in the current methods. Firstly, most methods^20, 21, 23, 26-29^ use a regression-based deep learning model, which emphasizes the learning of low-frequency mode^30^, resulting in the generation of excessively smoothed predictions. A few methods utilize CycleGAN-based approaches, which may be susceptible to unstable training processes and limited capability in handling geometric transformations^31^. Secondly, many current methods are designed for light microscopy^21, 25-27^, whereas EM images contain complex nanoscale structures, underexplored noise models, and single-channel limited image information, rendering image quality enhancement tasks more difficult. Third, none of the current methods can provide reliability assessments for their predictions. Since denoising and super-resolution are inherently ill-posed problems with multiple possible solutions, biologists may doubt the validity of the processed images without any uncertainty measures.

In this study, we propose EMDiffuse, a diffusion model-based package for EM applications, aiming to enhance EM ultrastructural imaging and expand the realm of vEM capabilities. Diffusion models have demonstrated superiority over regression-based models^32-34^ and exhibit greater stability in training than GAN-based models^35^ regarding image generation and restoration tasks due to their distinctive diffusion-based training schemes. Diffusion models also excel at preserving resolution and fine details, which is critical for imaging intricate nanoscale cellular structures with EM. Here, we adopted the diffusion model for EM applications and developed EMDiffuse-n for EM denoising, EMDiffuse-r for EM super-resolution, and vEMDiffuse-i and vEMDiffuse-a for generating isotropic resolution data from anisotropic volumes for vEM. Moreover, we demonstrate the self-assessment capability of EMDiffuse, taking advantage of the inherent generative capability of the diffusion model.

## Results

### EMDiffuse exhibited outstanding denoising performance

We developed EMDiffuse-n, the noise reduction pipeline of EMDiffuse, to denoise EM ultrastructural images and accelerate EM imaging (**Fig. 1a**). The EMDiffuse-n comprises three stages, which are data collection, image processing, and resolution preservation modeling (**Fig. 1a**, Methods). The first stage involved obtaining EM training pairs with different acquisition times (Extended Data Fig. 1a). Then, a hierarchical approach was employed to precisely align and register the noisy images with the reference image (Extended Data Fig. 1b, Methods). Finally, a diffusion model that preserves resolution, namely DiReP, (Extended Data Fig. 2, Methods), was trained to diminish the noise in well-aligned raw images and produce the denoised image with a difficulty-aware loss function ^36^. In the inference stage, multiple outputs are sampled, ensembled, and analyzed (**Fig. 1a**, Extended Data Fig. 3a, Methods) to improve the reliability of EMDiffuse-n (Methods, Extended Data Fig. 3b and 3c). Specifically, we optimized the prediction generation methods based on the assessment of FSIM^37^ (Extended Data Fig. 3b) and prediction resolution (Extended Data Fig. 3c) and used K outputs (K = 2 in our experiments) for each raw input and computed the mean of these outputs as the ultimate prediction (Methods, Extended Data Fig. 3). Of note, we assessed the predictions with FSIM instead of PSNR as PSNR prefers over-smoothed results (Extended Data Fig. 4).

**Fig. 1:**
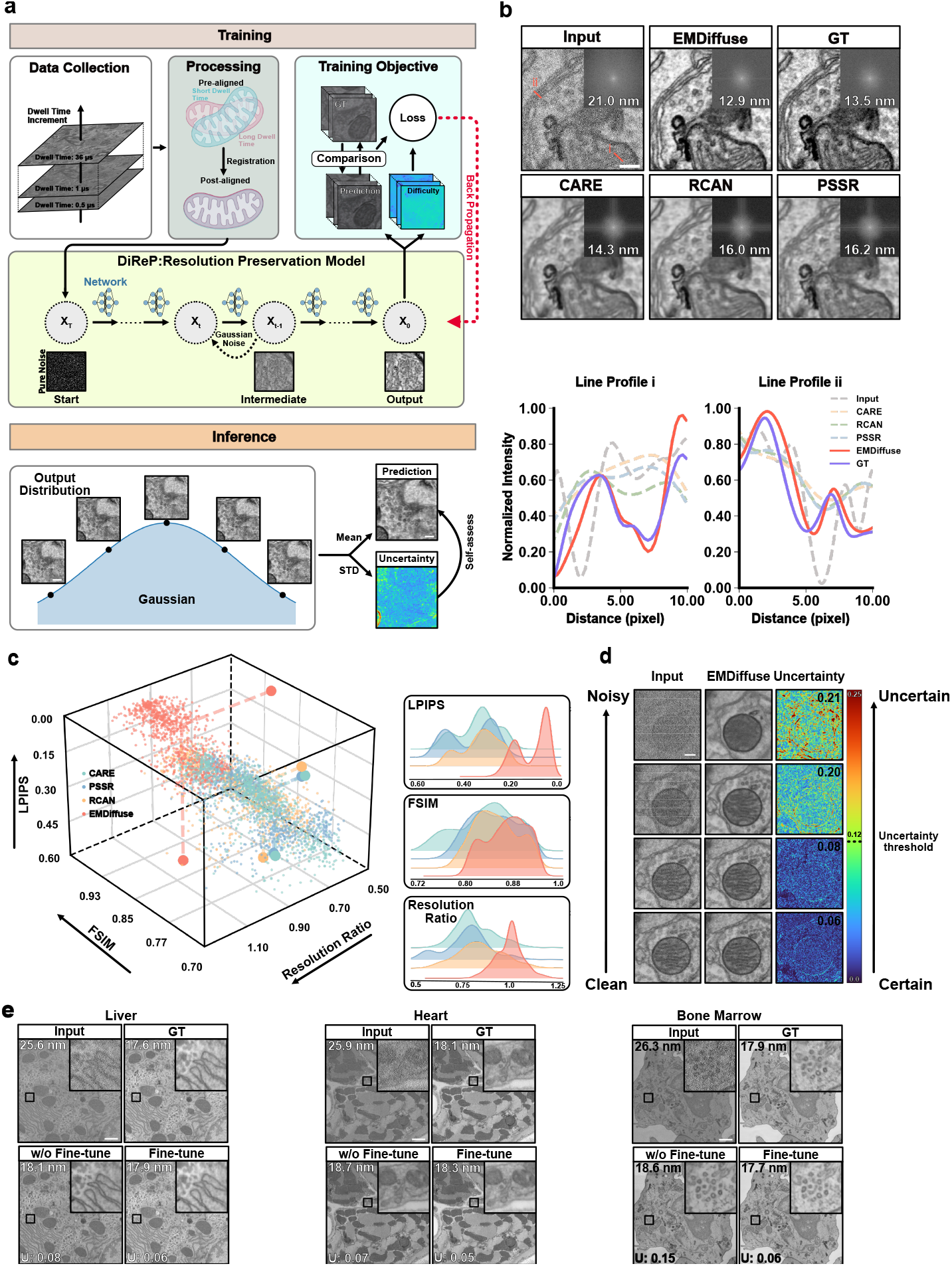
EMDiffuse exhibits excellent denoising capability and preserves image resolution for ultrastructural imaging. **a**. Schematic of EMDiffuse-n. In the training stage, paired data of various noise levels were collected and processed with a difficulty-aware training objective implemented. In the inference stage, EMDiffuse-n estimates and reduces the noise of raw EM images. **b**. EMDiffuse reduces the noise and enhances the resolution of EM images with a representative region of a mouse brain cortex tissue processed by EMDiffuse-n, alongside a comparison of the resolution and artifacts in the same image processed with denoising algorithms. GT: ground truth image; Input: input image, CARE, PSSR, and RCAN: generated images by CARE, PSSR, and RCAN algorithms from the input image, respectively. Top-right of each panel is the Fourier power spectrum. The resolution of processed images is shown on the Fourier transform image. Bottom panels are line profiles across the lines drawn in the Input panel. **c**. Quantitative performance assessment of images processed with EMDiffuse, CARE, PSSR, and RCAN algorithms using the Learned Perceptual Image Patch Similarity (LPIPS, lower values indicate superior performance) and the resolution ratio (ground truth resolution/prediction resolution, higher values indicate preservation of image resolution). Each point represents a unique test image of the mouse brain cortex, n = 960. **d**. Uncertainty map for EMDiffuse processed images at different noise levels enables self-assessment of uncertainty values of the prediction. Predictions with uncertainty values of the image below 0.12 (Methods) are counted as reliable outputs. **e**. EMDiffuse allows robust transferability. Representative input, EMDiffuse output without fine-tuning (w/o fine-tune), EMDiffuse fine-tuning with one training image (fine-tune), and GT images from the mouse liver, bone marrow, and heart tissues. A magnified region is positioned in the top right corner. The resolution of each panel is shown at the left top corner with nm as the unit. The uncertainty value of the results is shown in the bottom left corner. Scale bars, (**a, d, e**), 0.3 μm; (**b**), 0.1 μm.

Next, a comprehensive examination was conducted to evaluate the performance of EMDiffuse-n and other denoising methods (**Fig. 1b** and **1c**). EMDiffuse-n surpassed other denoising methods in preserving resolution and intricate ultrastructural information. This allows for adequately distinguishing mitochondria, cellular vesicles, endoplasmic reticulum (ER), and proximal plasma membranes (**Fig. 1b**, Extended Data Fig. 5 and Supplementary Video 1 and 2). Example line profiles (**Fig. 1b**) revealed that EMDiffuse-n successfully separated ultrastructure that is difficult to discriminate, such as mitochondria cristae and proximal plasma membrane, and produced results closest to the GT, while other denoising (*i*.*e*., CARE^28^, RCAN^38^, PSSR^23^) were incompetent in discerning these fine details. Of note, the Fourier power spectrum and resolution measurements^39^ indicated that EMDiffuse-n preserved the high resolution in the restored image (**Fig. 1b**). By contrast, other supervised deep learning models (*i*.*e*., CARE, PSSR, and RCAN) and self-supervised deep learning models (*i*.*e*., noise2noise^29^ and noise2void^19^) prioritized smoothness^40^, (white line across the center of power spectrum), and partially lost intricate structural details (**Fig. 1b** and Extended Data Fig. 5). In terms of quantitative evaluations, compared to other denoising methods, EMDiffuse-n demonstrated superior performance across three evaluation metrics, including LPIPS^41^, FSIM^37^, and resolution ratio, that assess models’ capability in generating accurate and high-resolution predictions (**Fig. 1c**). In particular, the resolution of predictions generated by EMDiffuse-n is high, closely matching the ground truth and exhibiting an approximately 30% improvement over predictions generated by other supervised deep learning models (**Fig. 1c**).

EMDiffuse-n addressed the common concern on the reliability of deep learning-generated data for scientific data processing (**Fig. 1d**). EMDiffuse-n was designed with the capability to reflect and self-assess the reliability of its predictions by calculating the standard deviation of K outputs for each raw image (**Fig. 1a**, Methods). EMDiffuse-n produced low uncertainty values when it was confident about its predictions (**Fig. 1d** and Extended Data Fig. 6). The overall uncertainty value of each input was calculated (**Fig. 1d**, Methods). We performed a study on a dataset with varying noise levels and identified an uncertainty value threshold, 0.12, that could assist biologists and microscopists in assessing the reliability of predictions (Methods). Predictions with higher uncertainty values, even though they may appear to be high-quality EM images, should be used with caution.

Finally, EMDiffuse-n achieved strong generalizability and transferability (**Fig. 1e**). Training the EMDiffuse-n model requires a substantial amount of paired data. Specifically, it requires 500 training pairs, which took one day to capture. To avoid extensive data-capturing for various EM ultrastructural imaging data, we explored the possibility of generalizing and transferring the pre-trained model to new types of biological samples (Extended Data Fig. 7a, Methods). We tested on datasets from three different tissues: mouse liver, heart, and bone marrow, and showed EMDiffuse-n had excellent generalizability and high transferability on images from biological samples different from training data (**Fig. 1e** and Supplementary Video 3). Moreover, we also showed that the performance of the pre-trained EMDiffuse-n model could be easily improved by fine-tuning the decoder with only a single data pair of 1768 × 1768 from new domains (Extended Data Fig. 7). After fine-tuning, the predictions from all three datasets showed low uncertainty values and high reliability (**Fig. 1e**). This opens the possibility of applying EMDiffuse-n in various EM application scenarios.

### EMDiffuse achieved superior super-resolution capability

We also applied EMDiffuse for the super-resolution task (EMDiffuse-r) that reconstructs a high-resolution image, which requires a long acquisition time, from a noisy low-resolution image, which needs only a short acquisition time (**Fig. 2a** and Extended Data Fig. 8). We optimized the prediction generation and computed the mean of two outputs for each raw input as the prediction (Extended Data Fig. 9a, Methods) and computed the uncertainty values for assessing reliability. Firstly, EMDiffuse-r has exhibited superior qualitative performance, particularly in improving resolution and resolving intricate details, while other tested methods introduced unwanted smoothness (**Fig. 2b**, Extended Data Fig. 9b Supplementary Video 4 and Video 5), which was consistent from low-resolution input images with different noise levels (Extended Data Fig. 9c). For example, EMDiffuse-r successfully discerned and separated the proximal synaptic vesicles (**Fig. 2b**) and mitochondria cristae (Extended Data Fig. 9c) while other methods failed to differentiate them. Then, the resolution values and Fourier power spectrum further confirmed that EMDiffuse-r did not introduce the undesirable smoothness and artifacts into predictions (**Fig. 2b** and Extended Data Fig. 9c). Quantitative comparisons in LPIPS, FSIM, and resolution ratio metrics (**Fig. 2c**) reveal that EMDiffuse-r outperformed other models in all three metrics and generated the most accurate and high-resolution predictions. Furthermore, the uncertainty value below the threshold indicated that the prediction was reliable except for extremely noisy raw inputs (**Fig. 2b** and Extended Data Fig. 9c). The generalizability and robust adaptability of EMDiffuse-r also enabled easy adaptation of the proposed model to a brand-new dataset with a single training data pair (**Fig. 2d** and Supplementary Video 6). Notably, by super-resolving a noisy 6-nm pixel size image into a clean 3-nm pixel size image, EMDiffuse-r doubled the image resolution and facilitated a 36× increase in EM imaging speed for the specific imaging setup used in this study.

**Fig. 2:**
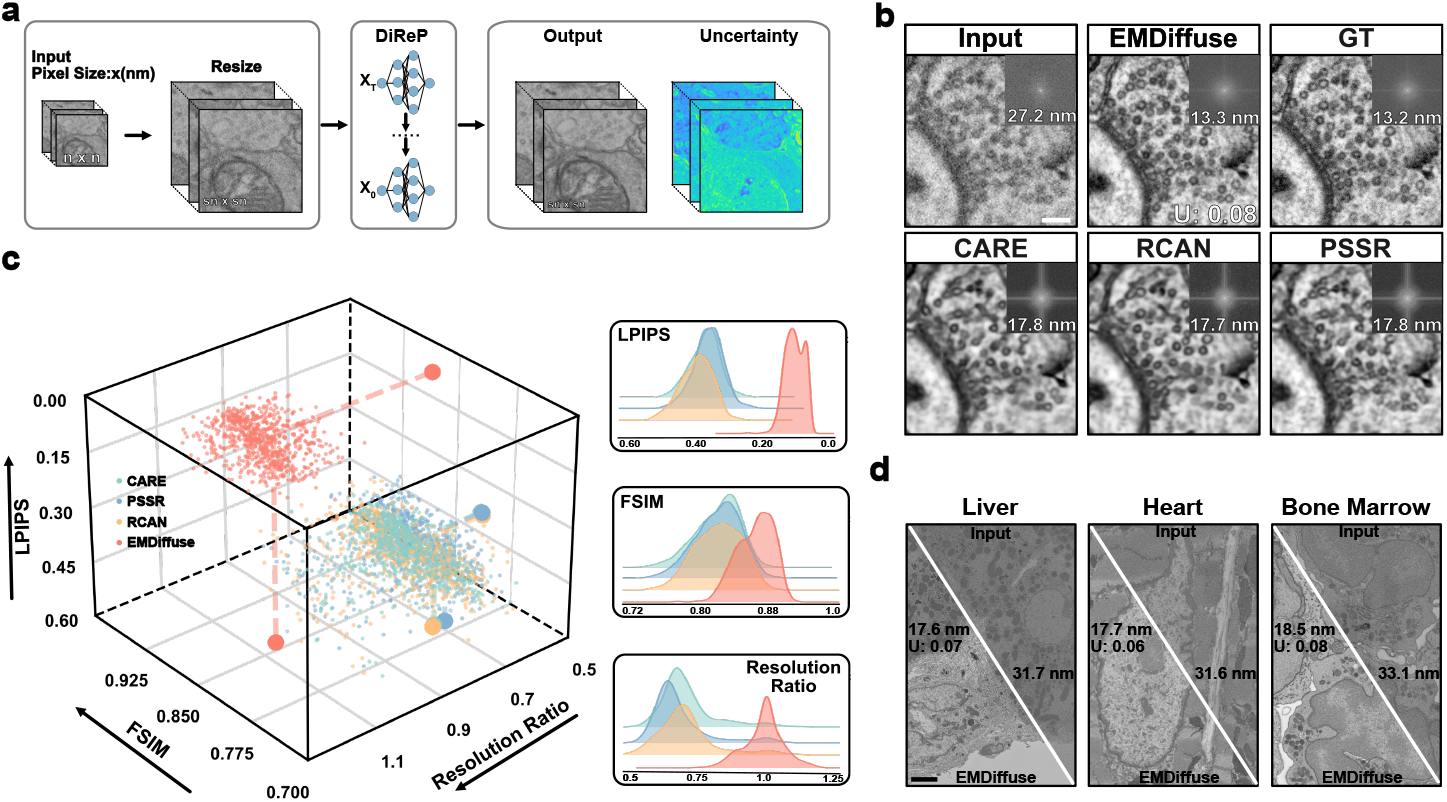
EMDiffuse achieves superior performance in super-resolution. **a**. Schematic of EMDiffuse-r for super-resolution of EM images. **b**. Representative region for comparing the super-resolution capability of EMDiffuse-r with CARE, PSSR, and RCAN. Top-right of each panel is the Fourier power spectrum. The resolution of processed images is shown on the Fourier transform image. The uncertainty value is shown in the bottom right corner. **c**. The 3D scatter plot and distribution plots show the quantitative performance assessment of EMDiffuse, CARE, PSSR, and RCAN for the super-resolution task. Each point represents a unique test image of the mouse brain cortex, n = 960. **D**. EMDiffuse for super-resolution is robust and transferable to other tissues, including the liver, heart, and bone marrow. Resolution values and uncertainty values are indicated on each panel. Scale bar, (**b**), 0.1 μm; **(d**), 0.3 μm.

### vEMDiffuse – EMDiffuse for enhancing volume electron microscopy

Isotropic resolution reconstruction from anisotropic vEM data can significantly accelerate vEM and expand its potential applications. In this work, we further extended EMDiffuse to vEM data and developed vEMDiffuse, which can generate isotropic volumes from anisotropic ones to reduce the number of layers to be captured for volume generation and accelerate vEM data acquisition (**Fig. 3** and **Fig. 4**). Specifically, we developed vEMDiffuse-i which incorporates a channel embedding (Methods) to generate layers between the 1^st^ and N^th^ layers by learning from a small volume of isotropic training data^13, 15, 42^ (**Fig. 3a**). In the inference phase, vEMDiffuse-i generates vEM data of isotropic resolution from an anisotropic one by generating intermediate layers between two consecutive layers of the anisotropic volume (**Fig. 3a**, Methods). Uncertainty values were calculated for predictions of each layer to ensure reliability. To validate the isotropic reconstruction capability of vEMDiffuse-i from an anisotropic volume, we downgraded a part of the isotropic volume (8 nm × 8 nm × 8 nm resolution) of the OpenOrganelle mouse liver (jrc_mus-liver)^43^ and OpenOrganelle mouse kidney (jrc_mus-kidney)^15^ to anisotropic volumes with 8 nm × 8 nm × 48 nm voxel size by removing the axial layers.

**Fig. 3:**
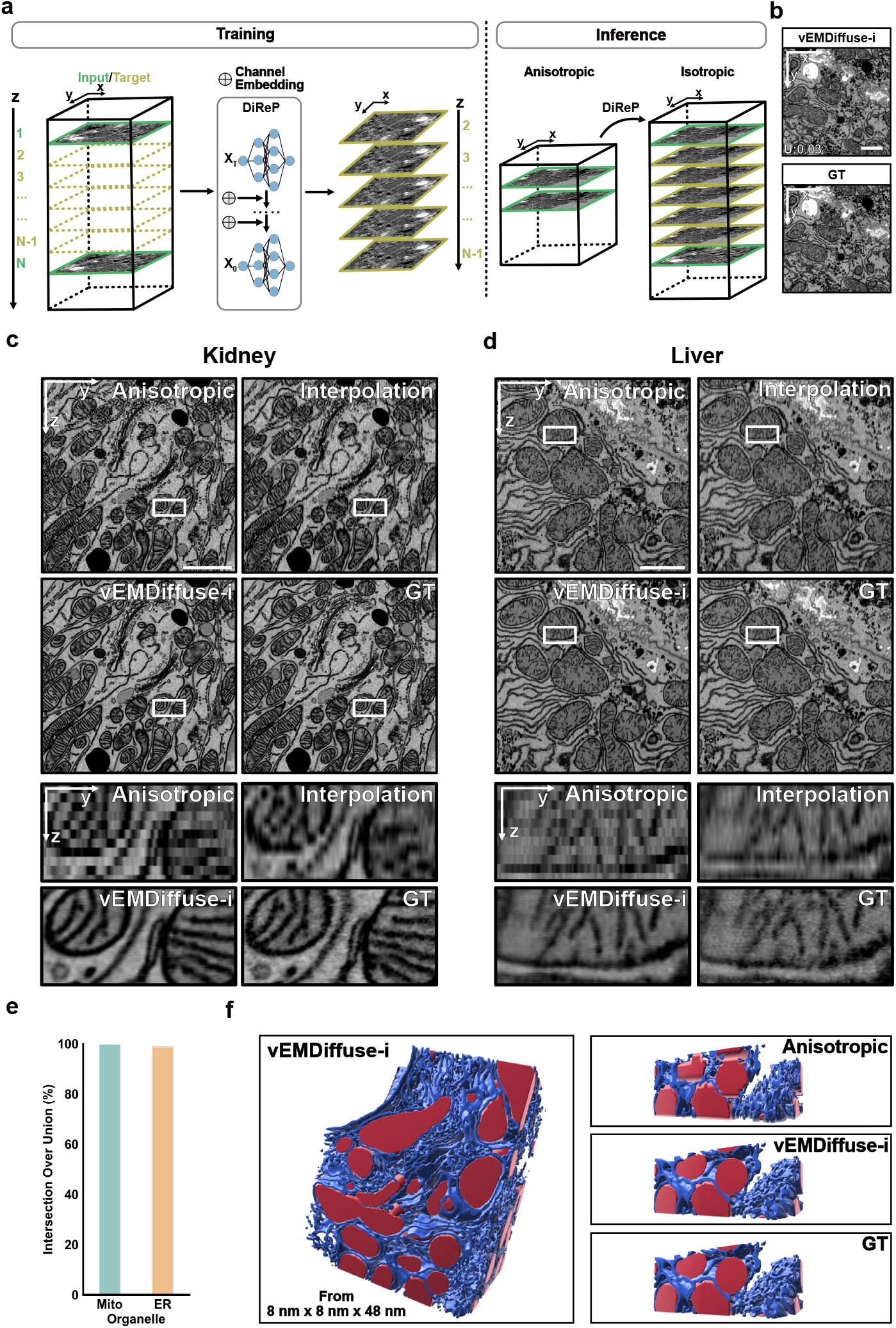
Accelerating isotropic volume electron microscopy (vEM) using vEMDiffuse for isotropic volume reconstruction from anisotropic data. **a**. Schematic of vEMDiffuse-i. vEMDiffuse-i learns to predict consecutive z-slices (*yellow*) with the preceding and following slices (*green*) as input. In the inference stage, vEMDiffuse-i achieves an isotropic resolution of vEM by generating intermediate layers between two layers of anisotropic volume. **b**. Representative generated XY view and the corresponding ground truth (GT) data. **c and d**. Representative vEMDiffuse-i generated YZ views of the Openorganelle mouse kidney dataset (jrc_mus-kidney) and the Openorganelle mouse liver dataset (jrc_mus-liver). Bottom panels are enlarged boxed regions of the anisotropic stack, interpolation stack, vEMDiffuse-i generated stack, and the ground truth (GT) isotropic stack. **e**. Segmentation mask overlap ratio on Openorganelle mouse liver dataset between the vEMDiffuse-i generated stack and the ground truth isotropic stack. **f**. Reconstructed 3D view based on the segmentation on vEMDiffuse-i generated stack (*left*) and comparison (*right*) of the 3D view of a zoomed-in volume between the anisotropic stack, vEMDiffuse-i generated stack and ground truth (GT) isotropic stack. Red, mitochondria; blue, endoplasmic reticulum (ER). Scale bar, (**b**), 2 μm; (**c, d**), 1 μm.

**Fig. 4:**
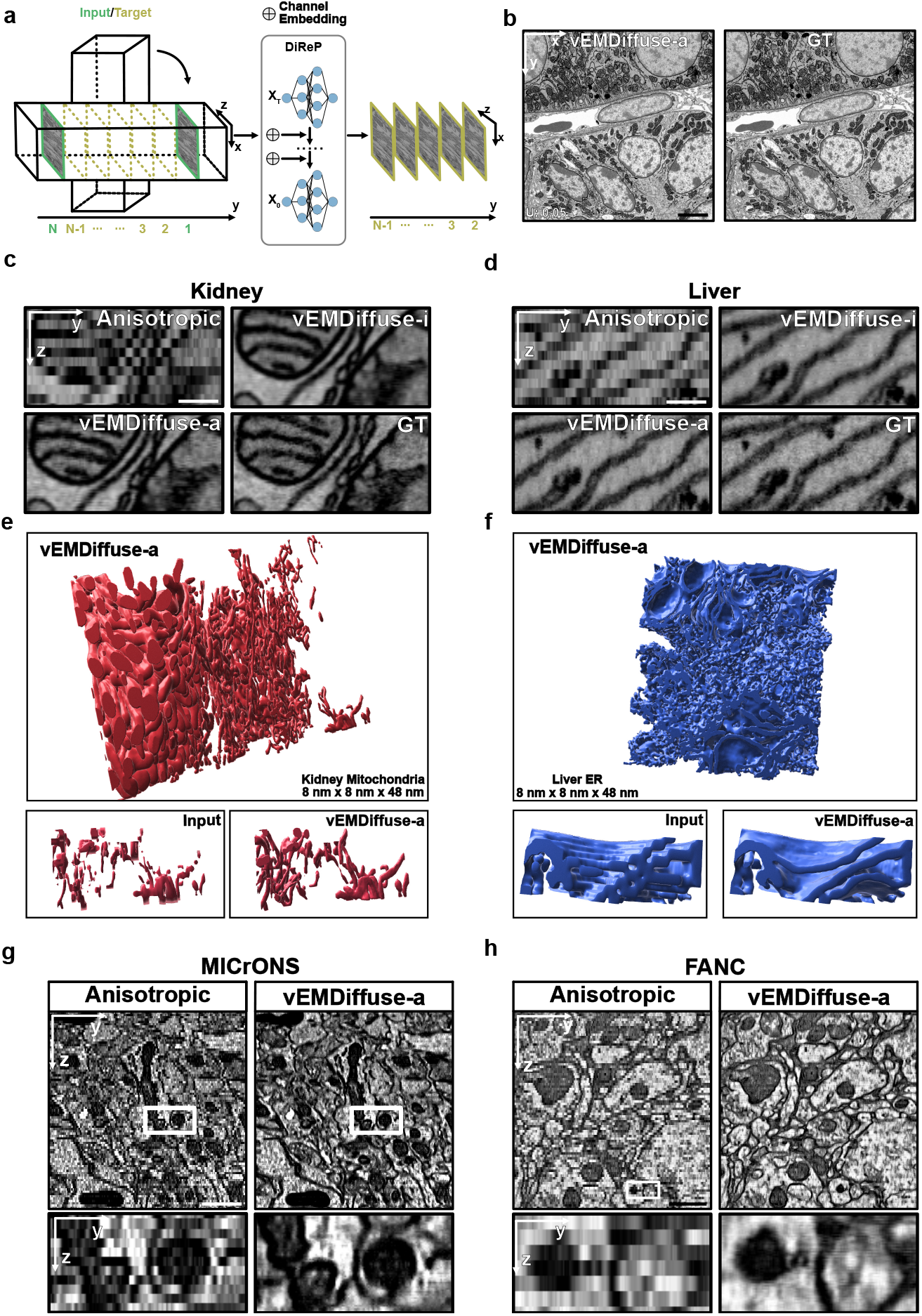
Expanding vEM realm with vEMDiffuse by generating isotropic resolution volumes with tissue array tomography type of data. **a**. Schematic of vEMDiffuse-a. During training, vEMDiffuse-a is trained to predict successive y slices (*yellow*) conditioned on the front and back y layers of anisotropic volume (*green*). For inference, same as vEMDiffuse-i, vEMDiffuse-a generates middle layers along the z-axis between two consecutive XY images from the anisotropic volume. **b**. Representative vEMDiffuse-a generated XY view of Openorganelle mouse kidney dataset (jrc_mus-kidney). **c** and **d**. Representative vEMDiffuse-a generated YZ view of Openorganelle mouse kidney dataset (jrc_mus-kidney, **c**) and Openorganelle mouse liver dataset (jrc_mus-liver, **d**). Shown are the anisotropic stack, vEMDiffuse-i reconstruction stack, vEMDiffuse-a reconstruction stack, and isotropic stack. **e** and **f**. 3D reconstruction of mitochondria and endoplasmic reticulum of the two datasets. Top panels are the reconstruction of the test volume. The bottom panels exhibit comparisons of an enlarged 3D reconstruction from the anisotropic stack and the vEMDiffuse-a generated stack. **g** and **h**. Representative vEMDiffuse-a generated YZ view (*top*) and enlarged region (*bottom*) of the MICrONS multi-area dataset and FANC dataset. Scale bar, (**b**), 1.5 μm; (**c, d**), 0.2 μm; (MICrONS in **g**), 0.8 μm; (FANC in **h**), 0.6 μm.

By reconstructing an anisotropic volume with n × n × sn resolution volume into an isotropic one with n × n × n resolution, vEMDiffuse-i expedited vEM imaging by a factor of s without compromising lateral resolution and axial continuity. First, the XY view example of the generated liver volume (**Fig. 3b**, Supplementary Video 7 and 8), and LPIPS, FSIM, and resolution metrics (Extended Data Fig. 10a) collectively indicated that vEMDiffuse-i could generate a volume with similar ultrastructural information and comparable axial resolution as the original isotropic volume. Then, an examination of a series of XY views showed that vEMDiffuse-i effectively maintained axial continuity and accurately and reliably (*i*.*e*., uncertainty values below the uncertainty threshold) replicated the ultrastructural changes of organelles within the isotropic volume, (Extended Data Figs. 10b, 11, and 12). Further, the YZ views (**Fig. 3c** and **Fig. 3d**) and XZ views (Extended Data Fig. 13) of the volume corroborated that vEMDiffuse-i accurately reconstructed the intricate structures of organelles, including mitochondria, ER and Golgi apparatus along the axial axis. By contrast, these ultrastructural details were lost in anisotropic volume and were not restored in ITK-cubic interpolated^44^ volume (**Fig. 3c, Fig. 3d**, and Extended Data Fig. 13). Moreover, we also applied vEMDiffuse-i on the OpenOrganelle T-cell^45^ (jrc_ctl-id8-2, Extended Data Figs. 14 and 15) and EPFL mouse brain datasets (Extended Data Fig. 16)^46^. vEMDiffuse-i consistently generated high-quality results with low uncertainty values and accurate organelle ultrastructure (Extended Data Figs. 14 and Extended Data Fig. 16a), suggesting its universal applicability on a variety of vEM datasets with a small volume of isotropic training data.

Segmentation of ultrastructural details (*e*.*g*., organelles) from vEM data is an essential task for vEM analysis, which often requires high-quality isotropic volumes^42^. Herein, we showed that vEMDiffuse-i predictions from anisotropic input volumes enabled accurate segmentation of organelles, achieving a similar quality as that using the isotropic resolution vEM captured with FIB-SEM as input (**Fig. 3e** and **Fig. 3f**). We trained an organelle segmentation model on the OpenOrganelle mouse liver dataset (Extended Data Fig. 17) and used it to segment the mitochondria and ER of vEMDiffuse-i generated volume ^42, 47^. The high intersection over union (IoU) score of the generated masks on the vEMDiffuse-i generated volume compared to the masks generated on the ground-truth isotropic resolution volume (**Fig. 3e**) indicated that organelle information generated by vEMDiffuse-i was precise in shape and position. The 3D rendering visualizations (**Fig. 3f** and Supplementary Video 9) demonstrated the accurate prediction of organelle ultrastructure in 3D, further demonstrating a high degree of reliability of the proposed model. In contrast, anisotropic stacks failed to reconstruct accurate 3D structures of organelles (*i*.*e*., mitochondria and ER), which represents a major limitation of almost all serial section-based vEM techniques.

### vEMDiffuse reconstructs isotropic resolution vEM volumes without isotropic resolution vEM training data

Acquiring high-quality isotropic vEM data from FIB-SEM is often impractical for most research laboratories, and the total volume that can be imaged in 3D by FIB-SEM is often small. To democratize access to isotropic vEM data and enable isotropic reconstruction of large tissue samples, we further developed vEMDiffuse-a which reconstructed isotropic volumes with only anisotropic training data (**Fig. 4a**). The division of input and target volumes for producing the training data in vEMDiffuse-a was the same as that of vEMDiffuse-i but occurred along the y-axis (as opposed to the z-axis in vEMDiffuse-i) (Methods). The inference stage was identical to vEMDiffuse-i, where vEMDiffuse-a generated intermediate layers between two consecutive XY layers of an anisotropic volume.

To consolidate our assumption that information in the lateral axis is transferable to the axial axis, we first applied vEMDiffuse-a to the manually downgraded anisotropic OpenOrganelle mouse kidney (jrc_mus-kidney) dataset^15^ and downgraded anisotropic OpenOrganelle mouse kidney (jrc_mus-liver) dataset, where some layers along the z-axis have been removed to simulate an anisotropic vEM data (Methods). First, although the XZ view training images contain undesired distortions (Extended Data Fig. 18a), we find that the vEMDiffuse-a model, given two consecutive XY view images as input, could produce intermediate layers with similar quality and resolution as the ground-truth ones (**Fig. 4b**, Extended Data Fig. 18b, Supplementary Video 10 and 11). The uncertainty value of generated layers also indicated that the predictions were reliable (Extended Data Fig. 18b). Further, the YZ and XZ views of the generated isotropic volume demonstrated that vEMDiffuse-a accurately captured the lateral information and leveraged it to enhance the axial resolution (**Fig. 4c, Fig. 4d**, Extended Data Fig. 19 and Supplementary Video 12). For example, the continuity of ER and contacts between ER and mitochondria (**Fig. 4c** and **Fig. 4d**) was accurately restored by vEMDiffuse-a. In contrast, in anisotropic volume, such information was lost, limiting investigations of organelle-organelle interactions in 3D. Moreover, the segmentation mask revealed that vEMDiffuse-a generated a volume that improved vEM organelle reconstruction, which will facilitate vEM investigation at the organelle level from an anisotropic volume (**Fig. 4e, Fig. 4f** and Supplementary Video 13). Again, anisotropic stacks failed to reconstruct accurate 3D structures of mitochondria and ER.

vEMDiffuse-a can be trained on any existing tissue array tomography type of anisotropic volumes and produce reliable isotropic volumes. We applied vEMDiffuse-a to two large open anisotropic tissue array tomography type datasets: the MICrONS multi-area dataset^48^ (**Fig. 4g**) and the FANC dataset^17^ (**Fig. 4h**). vEMDiffuse-a could generate an 8 nm × 8 nm × 8 nm voxel isotropic volume from the 8 nm × 8 nm × 40 nm resolution MICrONS multi-area volume (**Fig. 4g**, Extended Data Fig. 20) and a 4 nm × 4 nm × 4 nm voxel volume from the 4 × 4 × 40 nm FANC volume (**Fig. 4h**, Extended Data Fig. 21). The lateral image quality was comparable to the original volume (Supplementary Video 14 and 15) with uncertainty values below the threshold (Extended Data Fig. 20b, Extended Data Fig. 21b), while the axial resolution was boosted to an isotropic level, enabling researchers to observe seamless organelle ultrastructural changes along the z-axis with tissue array tomography data (**Fig. 4g, Fig. 4h** and Supplementary Video 16). By reconstructing an anisotropic volume into an isotropic one, vEMDiffuse-a enabled the visualization of cellular structures in 3D, paving the way for researchers to map the isotropic 3D ultrastructure of organelles within large tissue samples and pushing the boundaries of applications with vEM.

## Discussion

This study introduces EMDiffuse, a suite of deep learning-based methods that enhance the imaging power of ultrastructural imaging with EM and vEM. EMDiffuse demonstrates strong capabilities for expediting imaging processes and improving imaging quality through denoising and super-resolution tasks. More importantly, EMDiffuse ventures into an uncharted territory of isotropic vEM volume generation and broadens the horizons for vEM applications.

Leveraging a state-of-the-art image generation paradigm, the diffusion model, EMDiffuse exhibited superior performance in augmenting EM imaging while preserving the image resolution and nanoscale biological structure details. The superior performance of EMDiffuse can be attributed to the diffusion model’s ability to estimate data distribution gradients and gravitate towards regions with high data density^49^. This approach facilitates the learning of complex data distribution and detailed information, in contrast to regression-based models that primarily focus on learning the low-frequency information^30^. Moreover, EMDiffuse demonstrated remarkable robustness and transferability to facilitate effortless adaptation to any EM datasets with only one pair of fine-tuning images. Practically, microscopists may acquire a single training pair comprising a clean and noisy image of the target tissue to fine-tune the model. After fine-tuning the EMDiffuse, the rest images can be acquired at the high imaging speed required for noisy images. Of note, we observed the differences in the denoising performance for the direct application without fine-tuning, which likely represents the in-domain constraint of the deep learning model.

The use of artificial intelligence (AI) for image processing in biological imaging is on the rise. However, there is an understandable skepticism concern regarding the reliability of AI-generated images, as well as concern for potential artifacts in these images. EMDiffuse endeavored to address the reliability conundrum in bioimage processing deep learning applications with its self-assessment strategies. We also probed the upper bound of EMDiffuse’s reliable denoising and super-resolution capability with its self-assessment method. In our specific experiments, EMDiffuse can restore a high-quality image faithfully from a noisy image in the denoising tasks and reduce the data acquisition time by a factor of 18. Meanwhile, in the super-resolution task, EMDiffuse reduced the data acquisition time by a factor of 36 by restoring noisy and lower-resolution images to high-quality and higher resolution images. However, the expedition factor may vary among different EM instruments, depending on the instrument’s setup, particularly the detectors’ sensitivity. Nevertheless, it is possible to combine the acceleration capabilities of EMDiffuse and state-of-the-art EM instruments and detectors to push the limitations of attainable imaging areas of EM and enables the capturing of high-resolution images of larger areas–potentially more biological insights– within the same acquisition time.

By reconstructing an isotropic vEM stack from an anisotropic one, vEMDiffuse represents an inaugural effort to broaden the capabilities of vEM imaging. EMDiffuse-i and EMDiffuse-a were designed to tackle two types of vEM data from the most commonly used vEM techniques, FIB-SEM and serial section-based tomography methods, respectively. EMDiffuse-i demonstrated its robustness with isotropic reconstructions of four volumes from distinctive biological samples of various nanoscale structures and contrast (*i*.*e*., mouse brain, mouse kidney, mouse liver, and cultured T-cell and cancer cell). Compared with vEMDiffuse-i, which derives axial information from isotropic volume axial axes, vEMDiffuse-a transfers the knowledge acquired from the anisotropic volume lateral axes to the axial axes. It can be applied to any extant tissue array tomography-type anisotropic vEM data from serial section-based techniques without any additional data for training. Isotropic reconstruction and axial resolution enhancement on two public extensive vEM datasets–MICrONS multi-area and FANC–have exemplified vEMDiffuse-a’s capability to perform universal image transformations in existing large datasets, along with its potential impact on vEM and cell biology. Of note, we also expect that it is feasible for researchers to obtain a small volume of isotropic FIB-SEM data to train vEMDiffuse to reconstruct large isotropic volumes from serial section-based tomography methods of the same type of samples.

Recent investigations employing ultrastructural analysis with serial section-based tomography are largely limited to cellular scale segmentation and reconstruction^10, 12^. The advent of vEMDiffuse-a now enables faithfully isotropic reconstruction at the organelle scale for any serial section-based tomography vEM datasets, enhancing our ability to perform the three-dimensional reconstruction of numerous nanoscale organelles within tissues that were unfeasible to be investigated, hence, gaining a thorough comprehension of the intricate interactions between cellular organelles in 3D. Moreover, the capability of isotropic volume reconstruction from solely anisotropic datasets from serial section-based tomography methods is poised to democratize the isotropic vEM, which was only limited to researchers who have access to specialized FIB-SEM and enhanced FIB-SEM instruments. Ultimately, vEMDiffuse serves as the pioneering step in addressing the long-standing dogma in vEM techniques related to the trade-offs between isotropic resolution and attainable volume.

Further refinements to EMDiffuse are anticipated to fully harness the potential impact of this deep-learning model on biological imaging and cell biology. The stable diffusion model^33^ could be adopted as the backbone of EMDiffuse to accelerate the inference processes. The integration of algorithms of the EMDiffuse package can be investigated for further enhancements of EM imaging, particularly to accelerate the data acquisition and reconstruction of large isotropic volumes. The continued development of EMDiffuse will pave the way for tackling the grand challenge of reconstructing vEM volumes of multiple cubit millimeters, thereby facilitating connectomics mapping of larger animal models and enabling investigations of three-dimensional ultrastructural changes in large volumes of biological systems with isotropic resolution. Although the current study focuses on the processing of EM images, we envision that diffusion model-based deep learning methods hold substantial potential to advance a broader range of biological imaging techniques.

## Methods

### Sample processing and EM imaging

#### Mice and sample collections

C57BL/6 mice (Male, 10∼12-week-old) were provided by Animal Resources Centre (Australia), and the procedures were approved by the Animal Ethics Committee of The University of Western Australia. Following euthanasia, animals were perfused heparin (20U/mL) through the left ventricle, followed by a freshly prepared fixative solution containing 2% paraformaldehyde (PFA) and 2.5% glutaraldehyde in 0.1M sodium cacodylate buffer. Brain and liver were then collected and cut into small pieces (1-3 mm^3^), followed by the fixation at 4°C for 3 days. To image the bone marrow, mice tibias were collected and fixed at 4°C for 3 days prior to decalcification in 0.5M ethylenediaminetetraacetic acid (EDTA) for 3 weeks. Small bone fragments were cut for the following preparation.

#### Sample preparation

Tissues were processed using a modified protocol based on the serial blockface scanning electron microscopy protocol by the National Center for Microscopy and Imaging Research^50^. Briefly, following fixation, tissues were rinsed 5 times for 3 minutes each in 0.1 M sodium cacodylate before 1-hour incubation in 0.1 M sodium cacodylate buffer containing 2% osmium tetroxide and 1.5% potassium ferricyanide at 4°C. Samples were then rinsed and incubated with 1% thiocarbohydrazide for 20 minutes at room temperature. Next, samples were rinsed and incubated with 2% aqueous osmium tetroxide for 30 minutes at room temperature prior to the overnight incubation of 2% uranyl acetate at 4°C. The next day, samples were rinsed and dehydrated through a series of ethanol solutions at increasing concentrations (30%, 50%, 70%, 85%, 95%, 100%) for 10 minutes each, followed by two additional 10-minute incubations in 100% acetone. The samples were then infiltrated with Embed812 resin (Electron Microscopy Sciences) by incubating them in 33% resin (diluted in anhydrous acetone) for 2 hours, 66% resin overnight, and 100% resin overnight. Samples were embedded in fresh resin using polypropylene molds (Electron Microscopy Sciences) and polymerized in an oven for 48 hours at 65°C. After polymerization, samples were removed from the molds, block faces were trimmed, and 500-nm sections were cut using a Leica UC6 ultramicrotome.

#### Scanning electron microscopy

Sections of mouse tissues, measuring 500 nm in thickness, were mounted onto silicon wafers. Electron microscopy (EM) images were acquired using the backscattered electron detector using an FEI Verios scanning electron microscope (Thermo Fisher Scientific). For EMDiffuse-n, The dwell time of the electron dose on each point is changed to adjust the noise level of raw images. In the denoising task, each region is imaged 6 times with [0.5 µs, 1 µs, 1.5 µs, 2 µs, 4 µs, 36 µs] dwell time with 40,000× magnification.

### Denoise data pre-processing

The inherent distortions and horizontal-and-vertical drifts associated with EM imaging^51^ may result in misaligned training data (Extended Fig. 1a). Therefore, it is crucial to align the data properly for training the EMDiffuse model. In this study, we presented a two-stage registration strategy for coarse-to-fine alignment of the raw image and the ground-truth reference image. The coarse stage employed a key-point matching based registration technique^52^, which estimated a global homography matrix between them based on ORB (Oriented FAST and Rotated BRIEF) keypoints and feature descriptors^53^. The homography matrix was used to warp the raw image and aligned it with the reference image. However, the coarse stage could only align the two images globally, and local misalignments still existed (Extended Fig. 1b). Hence, in the fine stage, we employed optical flow estimation^54^ to estimate dense correspondences between pixels in two image patches which were further used to warp the raw image to achieve better local alignment with the reference image.

Additionally, for the sake of training efficiency, we cropped 256× 256 image patches with a stride of 196 to construct the data for training and inference. In the inference stage, for one raw image, the corresponding output patches from the model were stitched together using Imaris Stitcher (Oxford Instruments Group) to be the final output.

### EMDiffuse diffusion process and training objective

#### Denoising Diffusion Probabilistic Models in image quality enhancement

EMDiffuse’s fundamental basis is the Denoising Diffusion Probabilistic Model (DDPM)^32^, which belongs to a class of deep generative models. It can generate realistic data samples by reversing a diffusion process. The diffusion process, also known as the forward diffusion process, converts the target data distribution (*e*.*g*., the real image distribution) into a tractable probability distribution (*e*.*g*., the Gaussian distribution) by gradually adding noise to the data until they become pure noise. DDPM then learns to generate realistic samples from pure random noise by eliminating noise at each step, thereby reversing the aforementioned diffusion process. In the following section, we will elaborate further on the forward and reverse diffusion process.

#### Forward process

Given a sample **X**_o_ (e.g., a ground-truth reference image) drawn from the target data distribution *q*(**X**), the forward diffusion process, which gradually adds random gaussian noise with *T* discrete steps, produces a sequence of increasingly noisy samples **X**_1_, **X**_2_, …, **X**_*T*_ sampled from *q*(**X**_1:*T*_|x_o_).

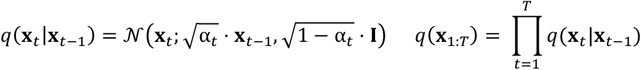

where **X**_*t*_ denotes the noisy data at step *t*, **I** is an identity matrix, and α_*t*_ ∈ (0,1) is the noise schedule parameter that controls the variance of noise at each step *t*. Specifically, the noise schedule is set so that **X**_*T*_ becomes pure random noise following a standard normal distribution. For this purpose, the noise schedule parameter : α_*t*_ initiates from a small value close to zero and grows progressively over time, reaching its maximum value at step *T*. For this purpose, we adopt the linear schedule as:

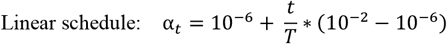

This process is equivalent to sample **X**_*t*_ at an arbitrary step *t* from *q*(**X**_*t*_|**X**_o_):

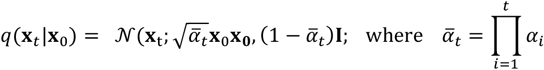

#### Reverse diffusion process

The reverse diffusion process uses a parametric model, i.e., a neural network, with parameters *θ*to invert the diffusion process and generates high-quality samples **x**_o_ from pure gaussian random noise **x**_*T*_. In our setting, we include the raw input image c_r_ (e.g., noisy input image) as a condition for the reverse diffusion process, which becomes a conditional process that can be expressed as below,

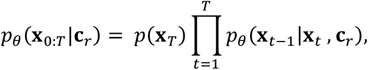

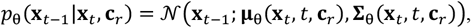

where **µ**_θ_ and **Σ**_θ_ are denoising functions realized by a deep neural network with parameters *θ* to predict the mean and covariance matrix and *t* is the reverse step index.

The reverse process is trained to maximize a variational lower bound, e.g., −*L*_*VLB*_(**x**_0_) of the log data likehood, i.e., log (*p*_θ_(**X**_0_)), which essentially targets at matching the joint distribution of the forward process, i.e., *q*(**X**_1:*T*_|**X**_0_), as:

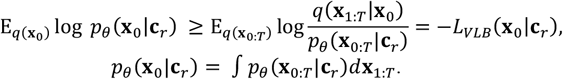

By minimizing *L*_*VLB*_(**x**_o_|**c**_r_), the network parameters *θ* are optimized.

Following DDPM^32^, we use a non-learnable fixed covariance matrix and adopt the reverse process parameterization of **μ**_θ_ as:

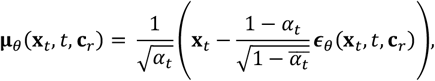

which uses a network ϵ_θ_ to predict the noise. DDPM empirically found that a simplification of *L*_*VLB*_(**x**_0_|**c**_r_) yield better sample quality in practice. We thus adopt the simplified the training objective:

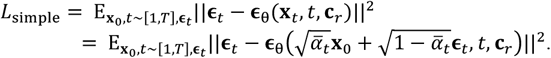

We observed that EMDiffuse training often got stuck at the hard cases (e.g., extremely noisy raw images in denoising task). Therefore, inspired by an existing method^36^, we added an additional prediction head that produced a difficulty assessment map **σ**_*i*,_ and used it to weight the prediction error. This resulted in a difficulty-aware loss function^36^:

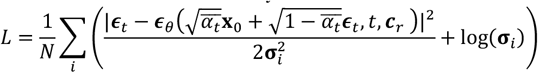

#### Inference stage

To generate samples from the target distribution, we start from a random noise sample *x*_*T*_ and iteratively apply the denoising function *p*_θ._(*x*_*t-1*_|*x*_*t*_) in reverse order, from step T to step 0:

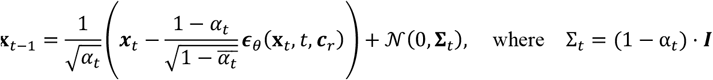

Also, each step is also guided Through step-by-step denoising, we can approximate the target distribution *p*(*x*_o_) in the final step *x*_o_ as the final prediction from the diffusion model. As for uncertainty prediction, we adopt the last step (i.e., t = 0) of the diffusion process as the final variance map.

### DiReP network architecture

#### U-Net architecture

In our approach, we employed a U-Net^47^ architecture for ϵ_θ_(**x**_*t*_, *t*, **c**_*r*_) akin to the one implemented in DDPM (Extended Data Fig. 2). U-Net is particularly well-suited for image denoising and super-resolution tasks, owing to its ability to efficiently capture both local and global contextual information from input images. The U-Net architecture consists of two main components: the encoder (Contracting Path) and the decoder (Expanding Path). These two elements are interconnected through skip connections, which facilitate the propagation of high-resolution spatial information from the encoder to the decoder. This feature enables U-Net to preserve fine-grained details in the resulting image, which is crucial for tasks such as image denoising and super-resolution.

#### Global attention

Drawing inspiration from SR3^55^, we incorporated a global attention layer, akin to the self-attention mechanism in transformers, into the U-Net architecture. This enhancement boosted model’s performance by allowing it to capture long-range dependencies and contextual information across the entire input image. Specifically, the global attention layer is placed at the bottleneck of the U-Net i.e. between the encoder (contracting path) and the decoder (expanding path), in order to reduce the computation costs.

A global attention layer operates as follows. Firstly, we use three linear layers to transform the input feature map, which is derived from the contracting path. This transformation produced the Query (*Q* ∈ *R*^*H*×*W*×*C*^), Key (*K*∈ *R*^*H*×*W*×*C*^) and Value (*V* ∈ *R*^*H*×*W*×*C*^) embeddings, where *H* and *W* represent the spatial height and width of the feature map, and *C* represents the feature dimension. Next, we compute the attention as:

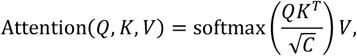

where the “sof*t*max” operation normalize the attention weights, resulting in a summation of 1. The attention module refines the feature of a single pixel location by softly aggregating the information from all spatial locations based on their similarities. This process enables the features to effectively capture global information.

### EMDiffuse-n training

In this work, the EMDiffuse networks were trained on a computer workstation equipped with four RTX 3090Ti NVIDIA graphic processing cards. The implementation was done using Python version 3.8 and PyTorch version 1.7.0^56^. The Adam optimizer^57^ with an initial learning rate of 5e-5 was employed to train EMDiffuse. The learning rate remained constant throughout the training processes since we observed no over-fitting except in transfer learning experiments. The checkpoint yielding the lowest reconstruction error on the validation dataset is used to test EMDiffuse on the test dataset. To enhance the robustness of EMDiffuse, data augmentation techniques such as flip, rotation, and Gaussian blur were applied during training. All the plots are generated using Matplotlib and Seaborn libraries in Python.

### Evaluation metrics

#### Noise level measurement

In the data acquisition process, the dwell time of the electron dose was adjusted to alter the noise level. However, using dwell time directly to represent noise level of raw images was not universally applicable across different microscopes and tissues. To quantitatively measure the noise level of each raw image, we employed a Blind/Reference-free Image Spatial Quality Evaluator (BRISQUE)^58^. This metric is designed to quantify an image’s perceptual quality of an image by examining the naturalness or statistical regularities present in the image. It can effectively evaluate the quality of images subjected to distortions such as compression, noise, or blurring, without requiring an original, undistorted reference image. We used this metric to evaluate the noise level of raw images during inference to optimize the prediction generation method.

#### PSNR

Peak Signal-to-Noise Ratio (PSNR) is a metric for measuring the quality of reconstructed or generated images with respect to their original ground-truth reference images^26, 28^. It is calculated by comparing the peak signal power (e.g. maximum image intensity) and noise power, measured by the mean square error between the generated image *I* and original reference image *R* as follows:

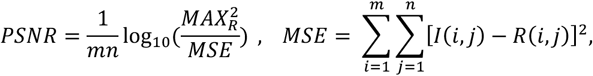

where *MAX*_*R*_ denotes the maximum pixel value of the ground-truth reference image *R*. A higher PSNR means better image quality. Although PSNR can quantify pixel-wise similarity between the generated image and the reference image, it lacks structural and perceptual awareness and is susceptible to being influenced by noise in the reference image. Areas with low PSNR successfully reconstruct the structures and deliver high-quality results compared to the ground truth reference image (Extended Data Fig. 5b). This is because the presence of unavoidable noise in the ground-truth EM image negatively influences the PSNR calculation. Besides, we find that PSNR favours blurry images, which has also been proven in^40, 41, 59^. However, blur-free high-quality structures are preferred in practical EM applications. Therefore, we use Feature Similarity Index Measure (FSIM)^37^ and Learned Perceptual Image Patch Similarity (LPIPS)^41^ which emphasize structural quality and align well with human judgment. Details of FSIM and LPIPS are discussed as follows.

#### FSIM

FSIM^37^ is a perceptual quality metric inspired by the human visual system. It assesses the similarity between images by examining their low-level features, such as phase congruency and gradient magnitude. It is calculated as follows:

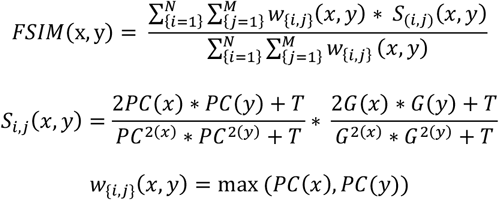

Where N and M are the height and width of the image respectively. *S*(*i, j*)(*x, y*) denotes the similarity measure between the phase congruency and gradient magnitude features of the pixels (i, j) in images x and y. *PC*(*x*) represents phase congruency calculated with the filter proposed ealier^60^. Phase congruency employs the phase of image Fourier transform to capture edges and other prominent structures in an image without being influenced by changes in lighting or contrast.

*G*(*x*) is the gradient magnitude calculated by gradient operators such as the Sobel operator^61^. Gradient magnitude, on the other hand, reflects local variations in intensity or colour within an image, offering insight into edges, textures, and intricate details. *w*_{*i,j*}_(*x, y*) denotes the perceptible significance of each pixel which is decided by the maximum value of phase congruency between two images. FSIM is specifically designed to capture the structural and textural information in images, which is a vital aspect of practical EM applications in biology. By concentrating on these features, FSIM delivers a more precise metric of structural similarity between images than PSNR^40, 59^.

#### LPIPS

LPIPS^41^ is a perceptual similarity metric that leverages deep neural networks. It is trained on an extensive dataset annotated by humans, which allows it to more precisely capture perceptual differences between images as perceived by human observers. LPIPS prioritizes the similarity of deep image features over pixel-wise differences, resulting in improved robustness and alignment with human perception. Consequently, LPIPS has become a widely adopted quality measurement metric in cutting-edge computer vision researches^33, 62^.

#### Resolution Ratio

In the context of electron microscopy, resolution refers to the ability of the microscope to distinguish between two closely spaced points or objects in the sample being examined. It measures the finest details that can be resolved in the sample’s structure. We further compute the resolution of input, generated image, and GT using image decorrelation analysis^39^. Then we computed the resolution ratio:

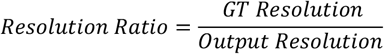

All the resolutions are transformed to nm as units by multiplying with pixel size.

### Optimization of prediction generation

When the noise level of a raw image is high, denoising and super-resolution tasks become notorious ill-posed one-to-many problems. In regions where noise dominates, multiple potential solutions exist (Extended Data Fig. 3a). To model such inherent ambiguities, we harnessed the power of diffusion models in generating diverse and multi-modal outputs. Specifically, given a single raw image **c**_*r*_, we sampled from the Gaussian distribution multiple times to produce inputs 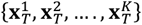 for the diffusion model, which in turn produces multiple results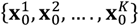. These diverse outputs captured the inherent multi-modal solutions. To alleviate the ambiguities in the predictions, we used the mean of these diverse outputs to be the final prediction **x**_0_.

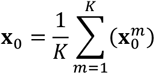

As the task becomes increasingly ill-posed with a higher noise level in the raw input image, we increased the number of *K* during inference. However, if the noise level of the raw image is low, incorporating a larger *K* to calculate the mean may introduce undesired smoothness into the prediction, potentially undermining the prediction resolution (Extended Data Figure 3b, 3c). Therefore, to achieve a balance of image quality and prediction robustness, we empirically determined the optimal number of K for a given input based on its noise level using the multi-noise-level mouse brain denoise dataset. We processed the data with different values of K and evaluated the quality of the outputs using FSIM as the metric (Extended Data Fig. 4a). We observed that K = 2 achieved superior FSIM scores in most cases and enhanced the predictions in most noise cases (noise level > 50). At extreme noise cases, using a larger K might contribute only marginal enhancement, and the resultant outputs might lack reliability, as the raw image is too noisy to restore some intricate structures (Extended Data Fig. 6). Users should be cautious about the predictions if the input is excessively noisy. This could also be reflected in our uncertainty score as described in the following. We measured the noise level of the raw image using BRISQUE which will be covered later.

### Assessing the reliability of predictions

#### Uncertainty map

Estimating the uncertainty of the model predictions is vital for enhancing reliability and aiding biologists in making informed decisions. We employed the variance of the K outputs from the fidelity enhancement module to supplement the model’s predictive uncertainty (Extended Data Fig. 6), as the prediction becomes increasingly trustworthy if multiple outputs agree, e.g., low variance. Thus, we calculated the standard derivation (STD) of K outputs 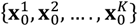 to supplement uncertainty estimation:

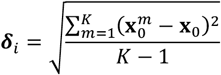

Given the uncertainty map, we obtained an overall uncertainty value for each input to help biologists to better determine the overall reliability of the prediction. In a practical setting, an inaccurate prediction of a small region or subcellular structure may lead to erroneous biological analysis and findings, and thus make the entire prediction image unreliable. We set a high constraint on the uncertainty value and extracted the uncertainty value by using the 99^th^ percentile of predictions in the uncertainty map. This means the whole prediction is deemed unreliable even if the model is unconfident about a small, predicted region or structure. We did not use the maximum value to avoid outliers. Biologists can compare the overall uncertainty value with the threshold introduced below to determine whether the prediction is reliable or should be used with caution.

#### Uncertainty threshold

We established an uncertainty threshold *τ* to guide biologists in practice. For one input, the prediction was considered as reliable if the overall prediction uncertainty was below *τ*. We determined the uncertainty threshold by examining the prediction accuracies of our multi-noise-level mouse brain denoise dataset using FSIM and LPIPS. We observed there is a transition valley between the low LIPIS (high FSIM) and high LIPSI (low FSIM) regions (Fig. 1c), which we found corresponds to images that are at the boundary of being accurate predictions (low LPIPS and high FSIM). We thus calculated the uncertainty of these samples and used the average of them to be the uncertainty threshold τwhich was found to be 0.12.

### EMDiffuse-n transfer learning

#### Transfer learning datasets

In transfer learning, we focused on three different tissues: mouse liver, heart, and bone marrow. The magnification rates of the three tissues were both 30,000×. The dwell time for raw images was set at 2 µs, while for ground truth images, it is 36 µs. These three datasets were imaged using the same SEM as the mouse brain cortex dataset. The same registration pipeline was applied to transfer learning datasets.

#### EMDiffuse-n data-efficient fine-tuning

First, we examined EMDiffuse-n’s generalization capability by directly applying the pre-trained model from the mouse brain cortex dataset to these new datasets (Fig. 1e). Second, we investigated the effectiveness of transferring the pre-trained EMDiffuse-n model to new tissues in a data-efficient manner by finetuning the model using few-shot tissue samples. To help the model avoid overfitting to the limited training samples and achieve better performance, we only finetuned the decoder and bottleneck layer parameters, as opposed to all model parameters (Extended Data Fig. 7a). We also investigated the performance of the model after fine-tuning using the different numbers of training samples (Extended Data Fig. 7b).

#### EMDiffuse-n fine-tuning details

During training, we used a learning rate of 1e-5 and fine-tuned EMDiffuse-n over 200 epochs on each dataset. All other configurations, such as the optimizer, remained consistent with those used in training EMDiffuse-n. To eliminate the impact of randomness, all experiments were replicated five times usingthe same training configurations.

### EMDiffuse-r for super-resolution

In the case of EMDiffuse-r, raw images were captured with a magnification factor of 20,000×, utilizing dwell time of 2 µs, 4 µs, and 6 µs. The ground-truth reference images were acquired at a 40,000× magnification factor and 36 µs dwell time, covering half the region size of the raw images. For data pre-processing, we rescaled the raw, low-resolution 128 × 128image patch to match its target high-resolution size of 256 × 256. This allows us to reuse the network architecture and pre-trained weights for the denoising task. We adopted the same training strategy as EMDiffuse-n and use the pre-trained model weights on the mouse brain cortex dataset to initialize the model. Similarly, we set parameter *K*to 2 for calculating the mean and uncertainty map as output (Extended Data Fig. 9a).

#### EMDiffuse-r transfer learning

To validate the transferability of EMDiffuse-r, we transferred EMDiffuse-r trained on mouse brain cortex dataset to mouse liver, mouse heart and mouse bone marrow datasets (the same datasets in EMDiffuse-n transfer learning). Since these three datasets were designed for transfer learning in the denoising task, the raw input was acquired with the same magnification rate as GT. Inspired by PSSR^23^, we addressed the scarcity of low-resolution images by manually downsampling the raw input with a short acquisition time. However, unlike PSSR, since we already had noisy images with low dwell time, we didn’t have to manually add Gaussian noise. Remarkably, we also achieved superior performance with a single pair of noisy downsampled images and GT (Fig. 2d) with the previously introduced transfer learning method.

### vEMDiffuse-i for isotropic reconstruction

vEMDiffuse-i was designed to restore an isotropic volume (voxel size 8 nm x 8 nm x 8 nm in our experiments) from an anisotropic one (voxel size 8 nm x 8 nm x 48 nm in our experiments) by interpolating intermediate layers 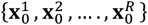 between two adjacent layers **c**^*u*^ (the upper layer), **c**^1^ (the lower layer) in an anisotropic volume (Fig. 3a).

#### Channel embedding

As axial resolution fluctuates across various datasets, satisfying the prerequisite for isotropic reconstruction requires varying the number of interpolation layers R. For example, within our experimental paradigm, vEMDiffuse-i is asked to interpolate 5 layers (R = 5) between **c**^*u*^ and **c**^1^ to enhance the axial resolution from 48 nm to 8 nm. However, for datasets with an axial resolution of 56 nm, vEMDiffuse-i should interpolate 6 layers (R = 6) between C^R^ and C^1^. Instead of modifying the model architecture (i.e., the output channel number of the last layer) to meet different R for different datasets, we introduced channel. The model of vEMDiffuse-i was fixed for any dataset, generating one layer at each iteration. The channel embedding takes the layer index as input, which indicates the relative distance between the target layer and upper layer **c**^*u*^, and embeds it into a feature vector using sinusoidal positional encoding^63^. For example, the number 3 indicates that vEMDiffuse-i is directed to generate the layer that is three layers distant from the upper layer **c**^*u*^ (4^th^ row in Extended Data Fig. 11). At the inference time, vEMDiffuse-i traverses from 1 to R (defined by the user depending on the dataset axial resolution) as the input of channel embedding, thereby guiding the diffusion model to sequentially generate intermediate layers. Since the model architecture is fixed, the pretrained model weights from other datasets can be transferred to a new dataset, reducing the training time by 48 hours.

#### vEMDiffuse-i training

During each training iteration, vEMDiffuse-i randomly selected one layer as the upper layer **c**^*u*^ in training isotropic volume and concatenated it with **c**^*u*+R+1^ layer of the volume (i.e., **c**^1^ = **c**^*u*+R+1^). The model was required to generate 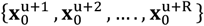 layers of the volume. The training objective of vEMDiffuse-i was:

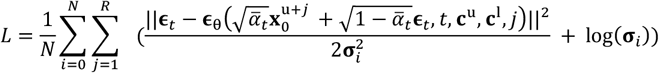

No overfitting was observed in our dataset, and we chose the checkpoint when vEMDiffuse-i’s performance stabilized on the validation set.

#### vEMDiffuse-i inference

During the inference stage, given an anisotropic volume {**c**^1^, **c**^2^, …., **c**^*M*^} with M layers, we traverse every pair of adjacent layers{**c**^*u*^, **c**^*u*+1^} of the volume: [{**c**^1^, **c**^2^}, {**c**^2^, **c**^3^}, … ., {**c**^M-1^, **c**^M^}]. For each pair of input, vEMDiffuse-i generated R images 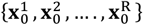 between them:

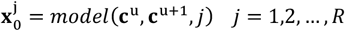

Then we inserted the generated layers between {**c**^*u*^, **c**^*u*+1^}. Thus, the final reconstructed volume 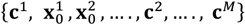 had ((M -1) × R + 2) layers. To enhancethe performance of vEMDiffuse-i, we obtained two outputs (K = 2) for each input and use their mean as the final prediction 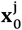.

#### vEMDiffuse-i inference dataset preparation

In the inference stage, we firstly cropped 1024 × 1024 × 1024 stacks (not used in the training stage) from OpenOrganelle Kidney isotropic volume^15^ and OpenOrganelle Liver isotropic volume^43^ as test datasets (voxel size is 8 nm x 8 nm x 8 nm). We manually removed some layers to simulate the anisotropic volumes. Starting from **c**^1^, we retained only the layers with indices whose remainder is 1 after divided by 6 while discarding the others:

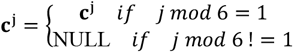

#### mod means the remainder after division

This reduces the axial voxel size from 8 nm to 48 nm.

### vEMDiffuse-a for isotropic reconstruction

In vEMDiffuse-i, we extracted layers along the z-axis of isotropic volume to train and thus we required isotropic volume for training. However, obtaining isotropic volume is impractical for many research groups. In vEMDiffuse-a, we trained the model’s isotropic reconstruction only with anisotropic volume.

#### vEMDiffuse-a model training

In vEMDiffuse-a, we extracted layers along the y-axis of the anisotropic volume (equivalent to the z-axis of isotropic volume in vEMDiffuse-i) to generate input and target pair to train (Fig. 4a). In the training stage, vEMDiffuse-a was required to generate 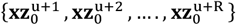 XZ layers of the volume given input pair of upper and lower XZ layer pair 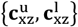 Of note, 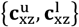 and 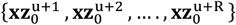 are both XZ view images exhibiting reduced image quality with line artifacts (Extended Data Fig. 20a and Extended Data Fig. 21a). We find the model is robust to these artifacts. We used the same training strategy as vEMDiffuse-i, but we noticed severe overfitting after a long training time, resulting in XZ-like predictions with clear line artifacts. To avoid overfitting, we terminated the training after 1200 epochs.

#### vEMDiffuse-a inference

The inference stage of vEMDiffuse-a was equivalent to that of vEMDiffuse-i. vEMDiffuse-a interpolated intermediate layers between upper and lower input pair 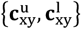. Again, of note, in the training stage, the input 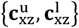 were XZ view images and the model was trained to interpolate layers along the y-axis. However, in the inference stage, the input 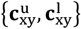 were XY view images of the anisotropic volume and vEMDiffuse-a interpolated layers along the z-aixs.

#### vEMDiffuse-a training and inference dataset preparation

To prove our concept that vEMDiffuse-a could transfer knowledge learned from the lateral axis (XZ views along the y-axis of anisotropic volumes at the training stage) to the axial axis (XY views along the z-axis of anisotropic volumes at inference stage), we firstly manually downgraded isotropic volumes: Openorganelle Kidney dataset^15^ and OpenOrganelle Liver isotropic volume^43^, and trained vEMDiffuse-a on the downgraded anisotropic volume. Given the isotropic volume 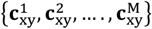, we reduced the axial resolution by removing layers:

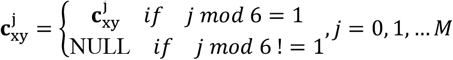

Of note, in vEMDiffuse-i, we only downgraded the volume in the inference stage while in the training stage, we still used isotropic volumes. Conversely, in vEMDiffuse-a, we downgraded the volume in both the training and inference stage. Then, we chopped along the y-axis to form training data: 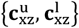 and 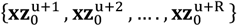 as elucidated above.

For original anisotropic datasets such as the MICrONS multi-area and FANC multi-area datasets, we directly sliced along the y-axis. For both MICrONS multi-area and FANC datasets, we reserved 2048 × 2048 × 128 stack as test datasets (not used in the training stage). In practice, researchers can also use the same anisotropic volume in the training and inference stage.

### Organelle segmentation

The organelle segmentation was trained with the isotropic volume and corresponding official segmentation masks provided by Openorganelle^15^. A 3D U-Net model^42^ served as the segmentation expert. We trained two separate models for mitochondria and ER segmentation using the bcedice loss function:

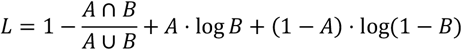

Where A represents the predicted segmentation mask of the model, and B denotes the GT mask. ∩symbolizes the intersection of two masks, while ∪represents the union. The segmentation model was trained using the Adam optimizer with an initial learning rate of 2e-4. We used small 170 × 170 × 80 patches with a stride size of 90 × 90 × 40 for both training and inference due to GPU memory limitations. In the inference stage, we classified pixels with a probability above 0.5 as organelle and the rest as background. We applied erosion and dilation post-processing techniques to remove small false positive regions and fill in small hollow holes. We rendered the 3D visualization with Imaris (Oxford Instruments Group), which also automatically filtered out some small regions.

## Acknowledgments

This work was supported by the Research Grants Council of Hong Kong (17102722, 17202422, 27209621) and the Australian Research Council (LP190100433). The work was conducted in the JC STEM Lab of Molecular Imaging, funded by The Hong Kong Jockey Club Charities Trust. We also thank Dr. Jing Guo for her assistance in image processing.

## Author Contributions

HJ, XQ and CL designed the experiments and wrote the paper. CL, KC, HQ, XC, and GC performed, collected, and assembled the experiments. HJ and XQ secured funding. All have commented on and edited the manuscript.

## Ethics declarations

The authors have declared that no conflict of interest exists.

## Data availability

Denoising and super-resolution training and test data for EMDiffuse are available at https://zenodo.org/record/8136295. For vEM, all OpenOrganelle datasets are downloaded from the OpenOrganelle website (https://openorganelle.janelia.org). The Openorganelle Kidney dataset is available at https://doi.org/10.25378/janelia.16913035.v1. The Openorganelle Liver dataset is available at https://doi.org/10.25378/janelia.16913047.v1. The Openorganelle T-Cell dataset is available at https://doi.org/10.6084/m9.figshare.14447541.v1. The EPFL mouse brain dataset is available at https://www.epfl.ch/labs/cvlab/data/data-em/. The MICrONS multi-area dataset can be downloaded from https://www.microns-explorer.org/. The FANC dataset can be downloaded from https://bossdb.org/project/phelps_hildebrand_graham2021. We also created a website for displaying high quality images and videos: https://www.haibojianglab.com/emdiffuse.

## Code availability

The source codes of EMDiffuse, several representative pre-trained models as well as some example images for testing are publicly accessible via https://github.com/Luchixiang/EMDiffuse.

## Extended data

**Extended Data Fig. 1.**
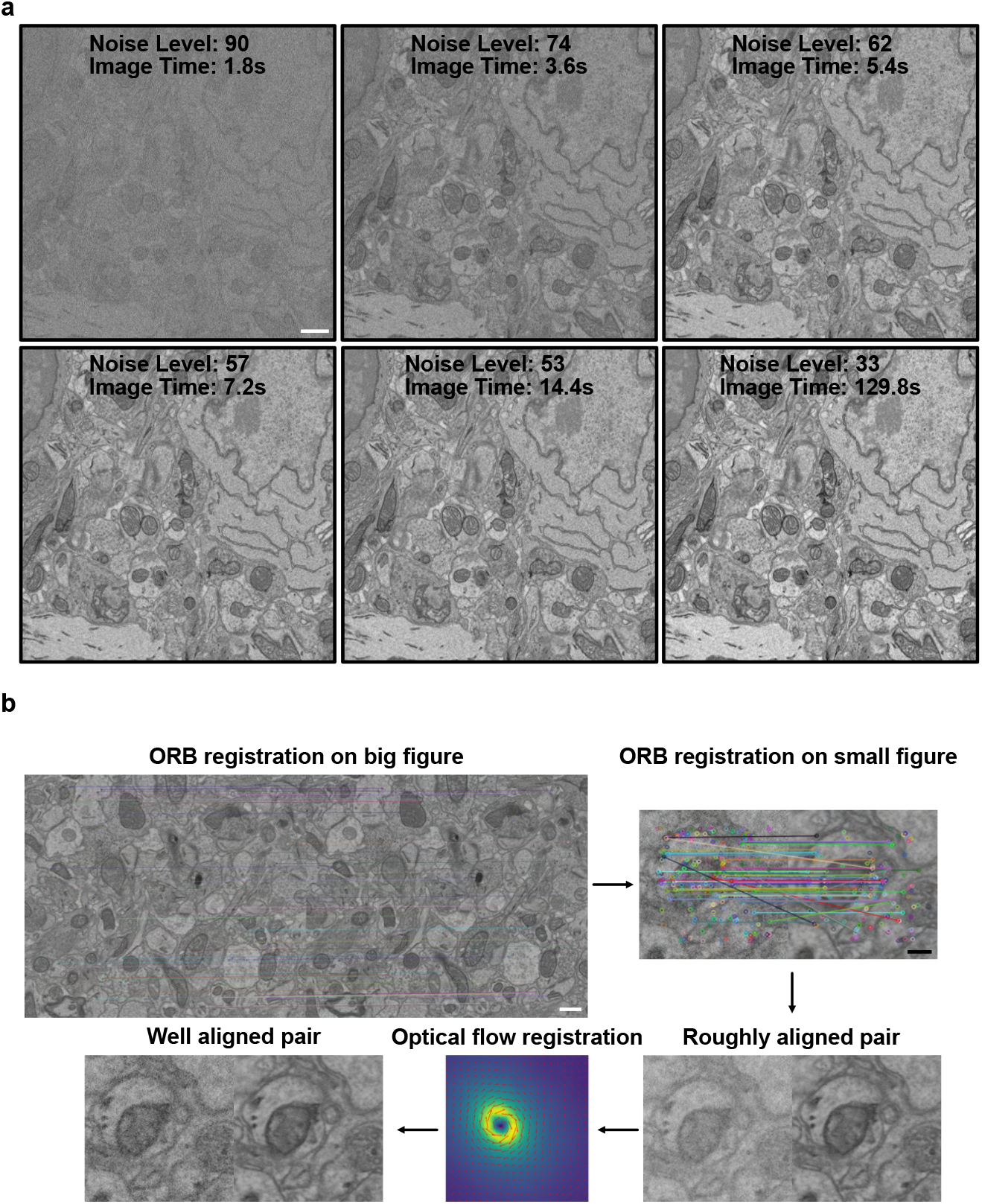
Denoise dataset acquisition and processing. **a**. Data acquisition. A dataset of electron microscopy images of the mouse brain cortex was acquired to train EMDiffuse. Various dwell times, ranging from 0.5 µs to 36 µs, representing different noise levels, are obtained to train a noise level-agnostic model and assesses the fidelity and limitation of EMDiffuse. **b**. Data processing. ORB registration is employed for the alignment of images of various noise levels. An additional optical flow registration is incorporated to enhance the alignment of training samples to mitigate the drift and distortion during the EM imaging for effective model training. Scale bar, white, 0.5 μm; black, 0.1 μm.

**Extended Data Fig. 2.**
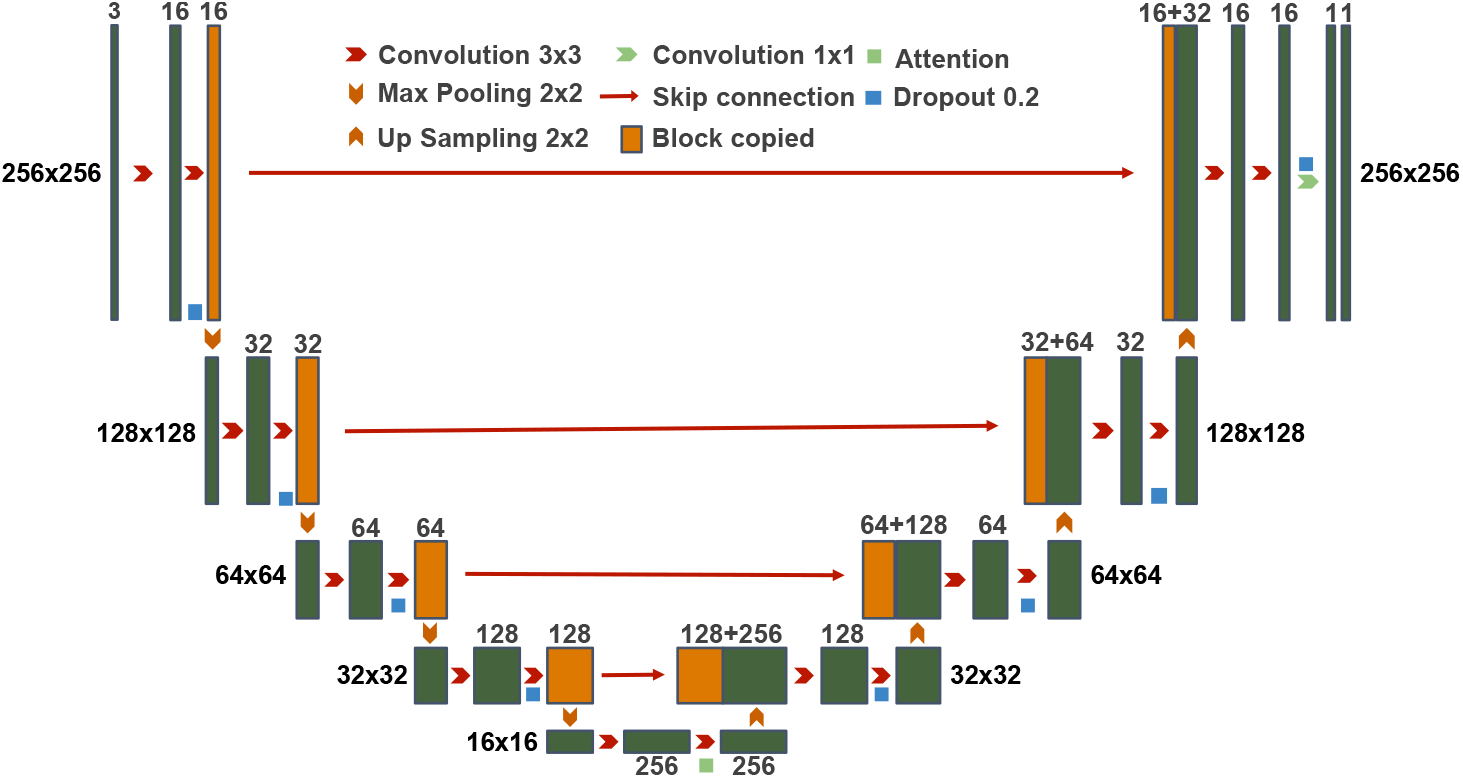
Network architecture of the DiReP model. A traditional U-shape model comprising an encoder and a decoder is employed in every step of the diffusion chain. Attention is added to the bottleneck of the model to improve model performance.

**Extended Data Fig. 3.**
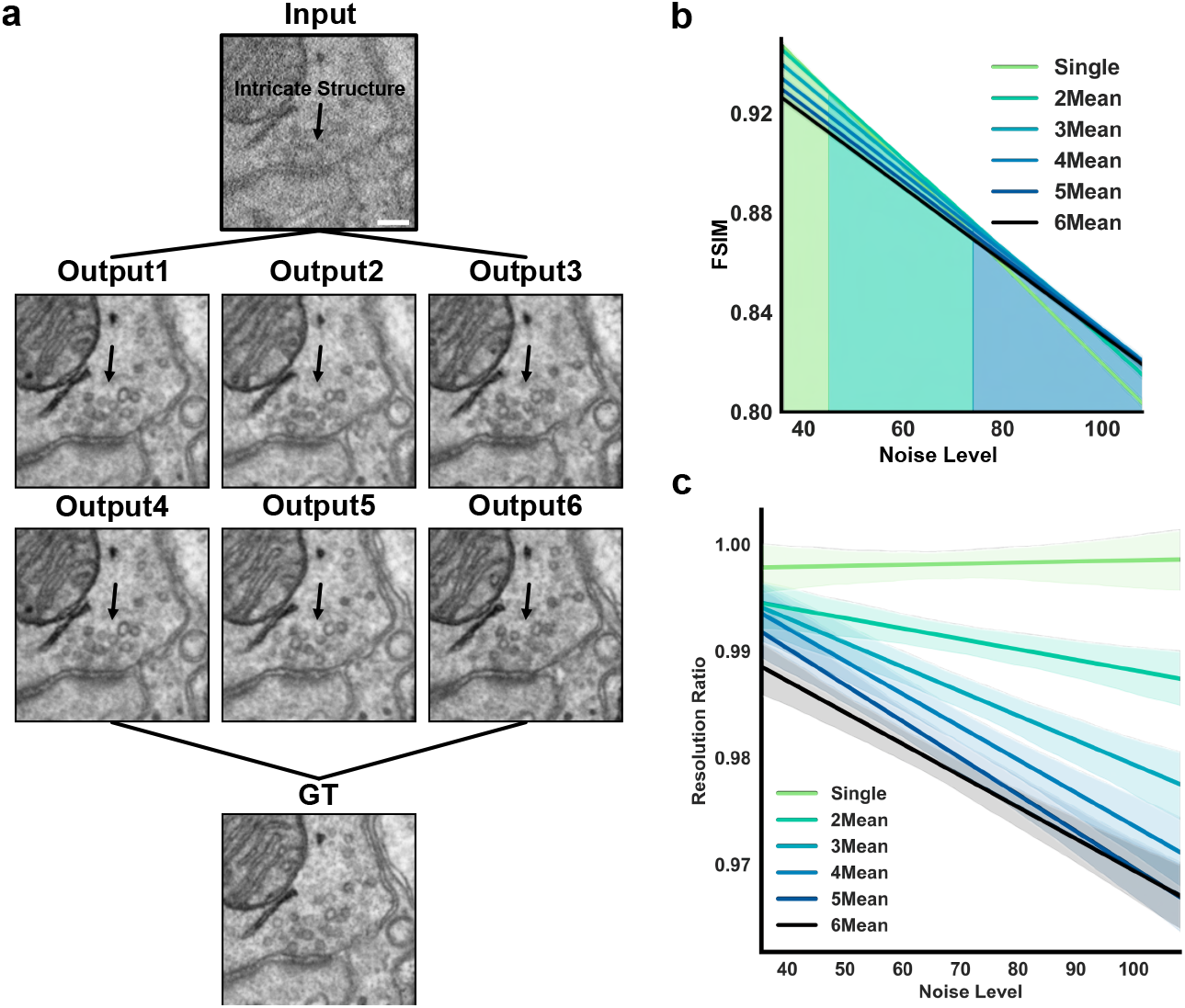
Optimization of prediction generation methods using multiple outputs enhances reliability while reserving resolution. **a**. EMDiffuse can generate multiple outputs, and averaging multiple outputs reduces the noise and improves the accuracy of the final prediction. In cases where the intricate structure is heavily dominated by noise, EMDiffuse can adopt the mean result of different testing times prediction as the ultimate prediction to increase fidelity. Scale bar, 0.1 μm. **b**. Assessments of EMDiffuse performance based on FSIM. Shown is the FSIM of EMDiffuse-processed images of different noise levels and the mean of different numbers of outputs. The number of outputs for generating the mean result that yields the highest FSIM is selected for EMDiffuse prediction. **c**. Quantification of resolution ratio (resolution of GT/resolution of EMDiffuse prediction) from the mean of different numbers of EMDiffuse outputs and different input noise levels. Scale bar, 0.1 μm.

**Extended Data Fig. 4.**
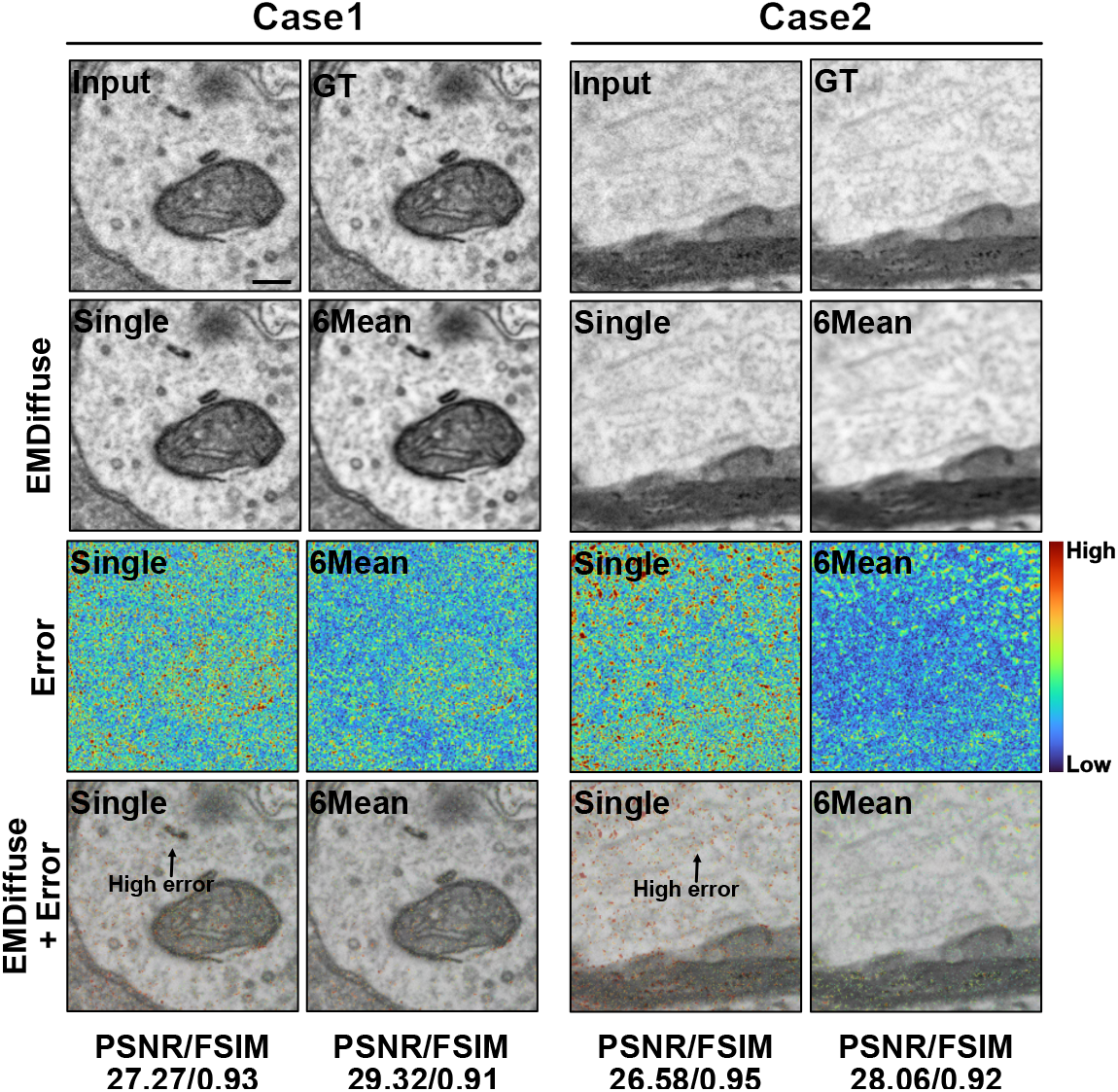
PSNR is not aligned with human perception. Illustration of PSNR drawback. The first row is the low noise level input and GT. The second row is the EMDiffuse prediction with the mean of six outputs (6mean) and without mean (single). Error maps (third row) and blend images of predictions and error maps (fourth row) indicating the PSNR score is significantly affected by noise and does not correlate with structural similarity and human perception. Scale bar, 0.1 μm.

**Extended Data Fig. 5.**
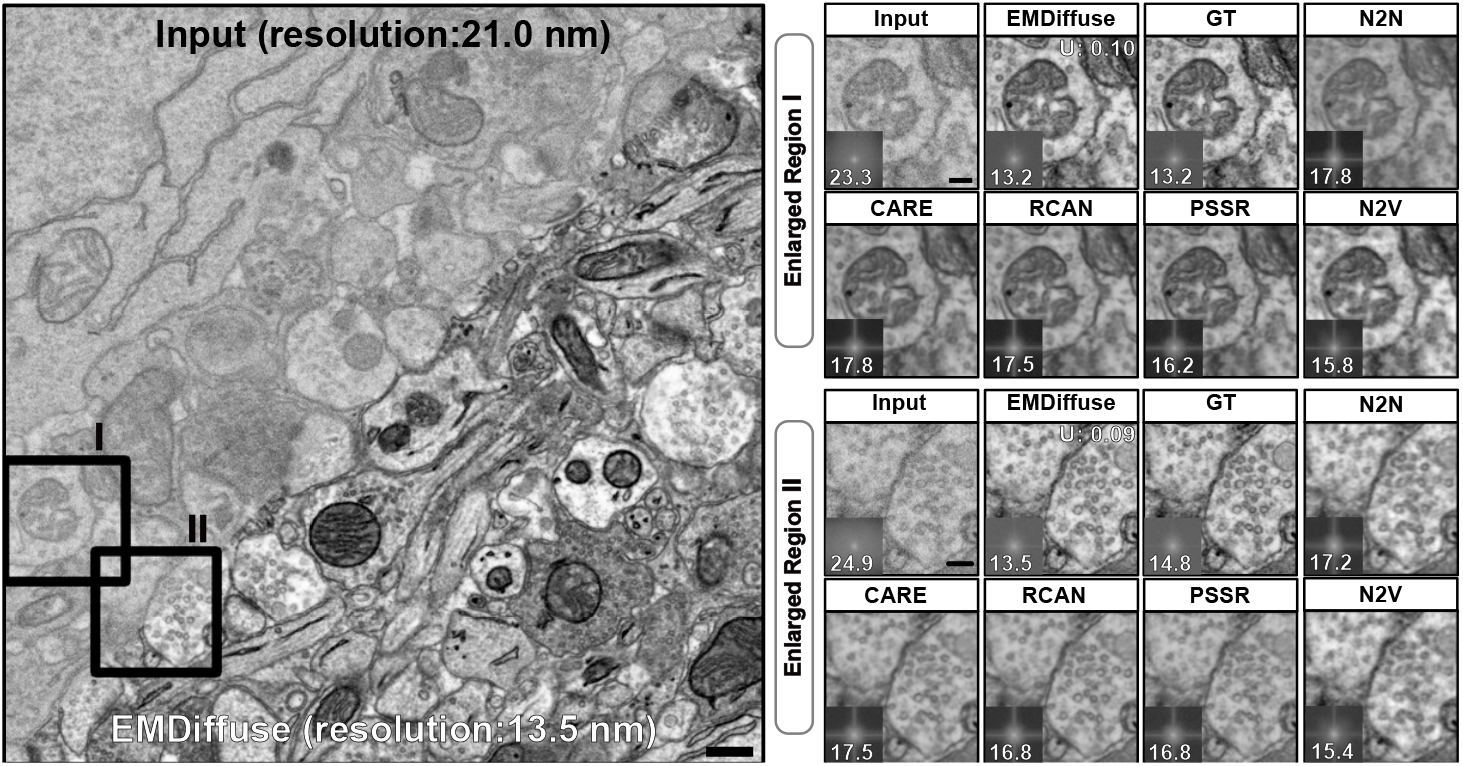
Additional examples of EMDiffuse-n for denoising EM images and comparison with other denoising methods. Additional examples of denoising of mouse cortex EM images compared to other denoising methods. Left: a denoising example containing both the noisy input and the EMDiffuse-n prediction. Right: two enlarged regions of raw input, ground truth (GT), and predictions from EMDiffuse, CARE, RCAN, PSSR, Noise2Noise (N2N), and Noise2Void (N2V) denoising methods. Resolution values and Fourier power spectrum are in each panel’s bottom left corner, showing that EMDiffuse preserves the resolution similar to the GT image and effectively reduces noise without undesirable over-smoothness. The uncertainty value is displayed in the top right corner. Scale bar, left, 0.3 μm; right, 0.1 μm.

**Extended Data Fig. 6.**
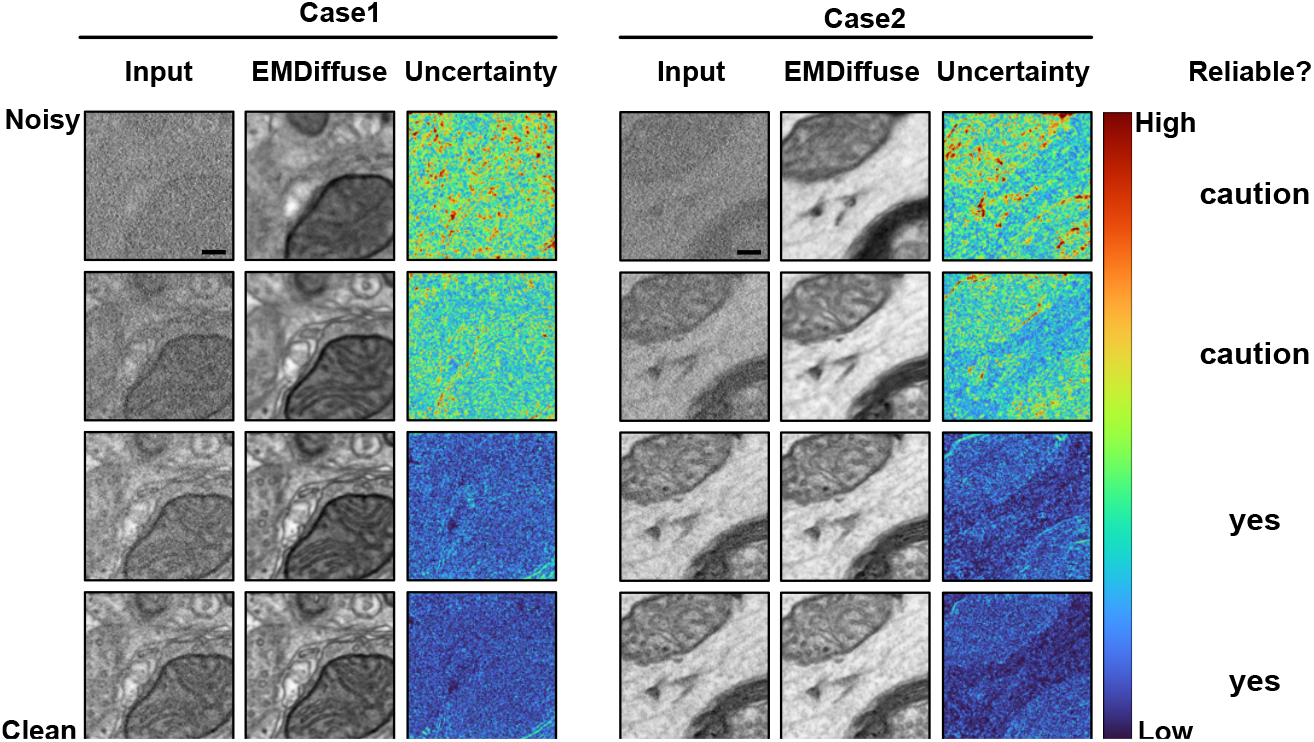
Uncertainty map for self-assessment of prediction reliability. EMDiffuse denoising predictions and uncertainty maps of two example regions with different noise levels. Each row consists of the noise levels, input image, EMDiffuse prediction, and the uncertainty map. A threshold of 0.12 for the total uncertainty is set for assessing the reliability of the final prediction, as shown in the last column. The caution signifies that researchers should use the prediction with caution while yes denotes that the prediction is reliable. Scale bar, 0.1 μm.

**Extended Data Fig. 7.**
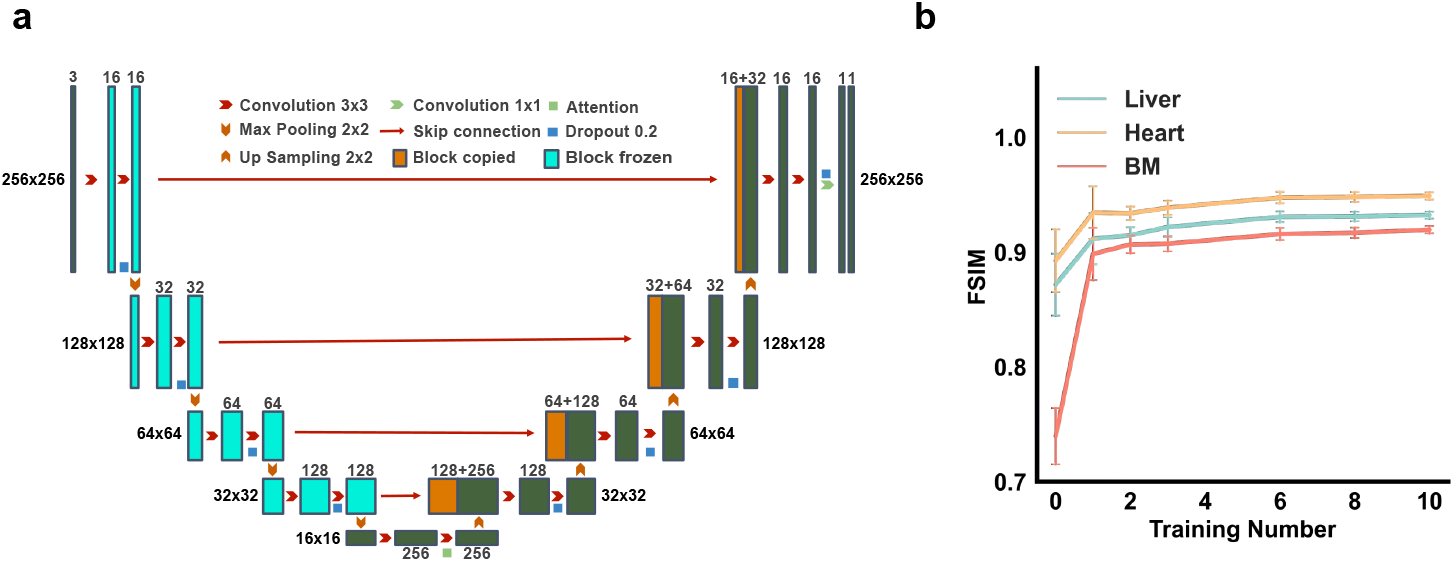
Encoder-frozen transfer learning for EMDiffuse enables fine-tuning with one training pair. **a**. Model architecture for EMDiffuse transfer learning. The encoder part of the model is frozen to prevent over-fitting and reduce the requirement for training data. The bottleneck with attention mechanisms is not frozen. **b**. FSIM results of EMDiffuse denoising performance using different numbers of transfer learning training images. The proposed half-frozen training approach achieved similar performance using only one training image across EM images from three different tissues, overcoming the limitations of a small fine-tuning dataset.

**Extended Data Fig. 8.**
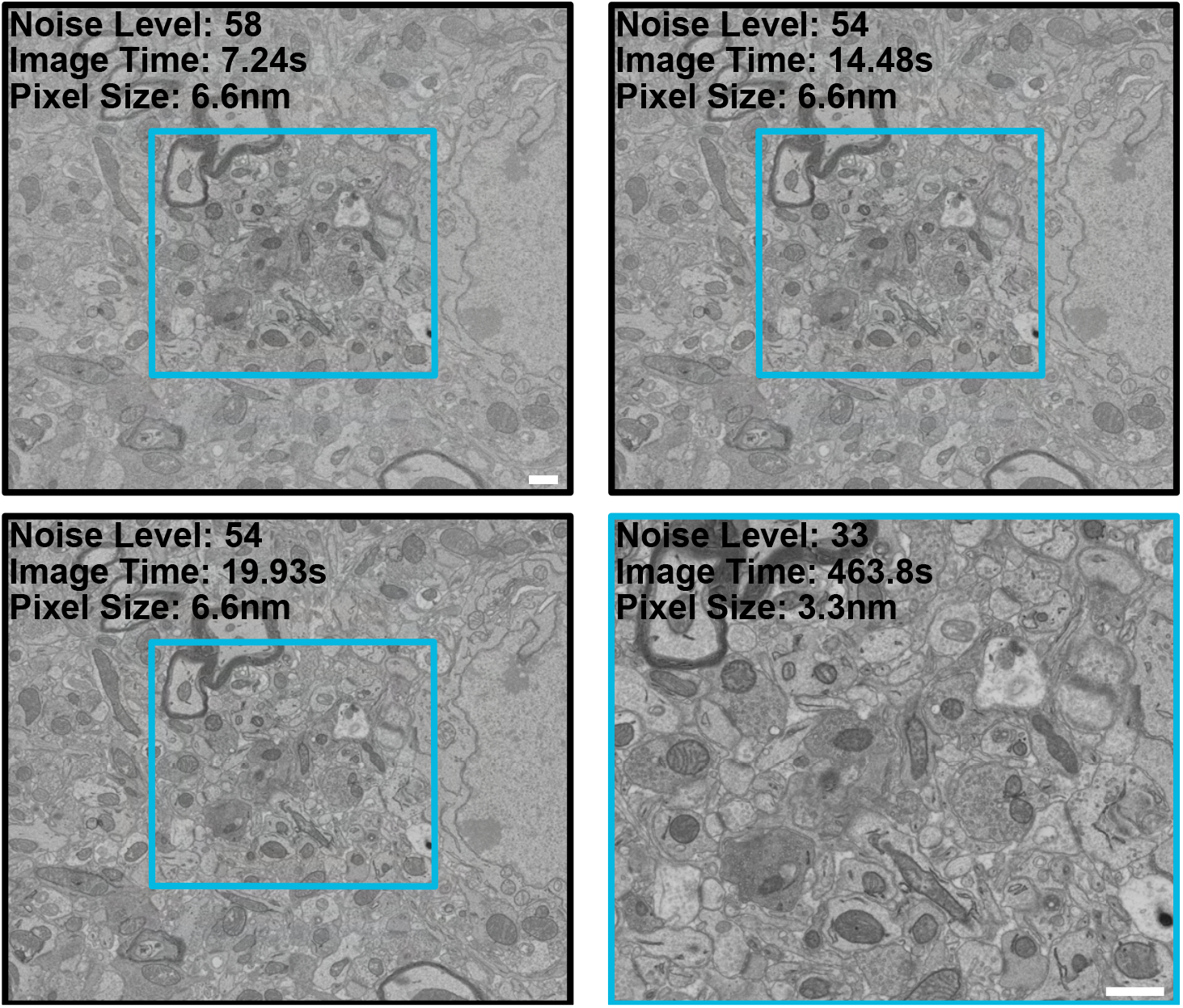
Super-resolution dataset acquisition. Examples of the source EM images are obtained with a pixel size of 6.6 nm (magnification rate of 20,000x), and the target image is obtained with a pixel size of 3.3 nm (magnification rate of 40,000x) at the central area in the source image. The dataset comprises three dwell times of input images ranging from 2 μs to 6 μs. Scale bar, 0.5 μm.

**Extended Data Fig. 9.**
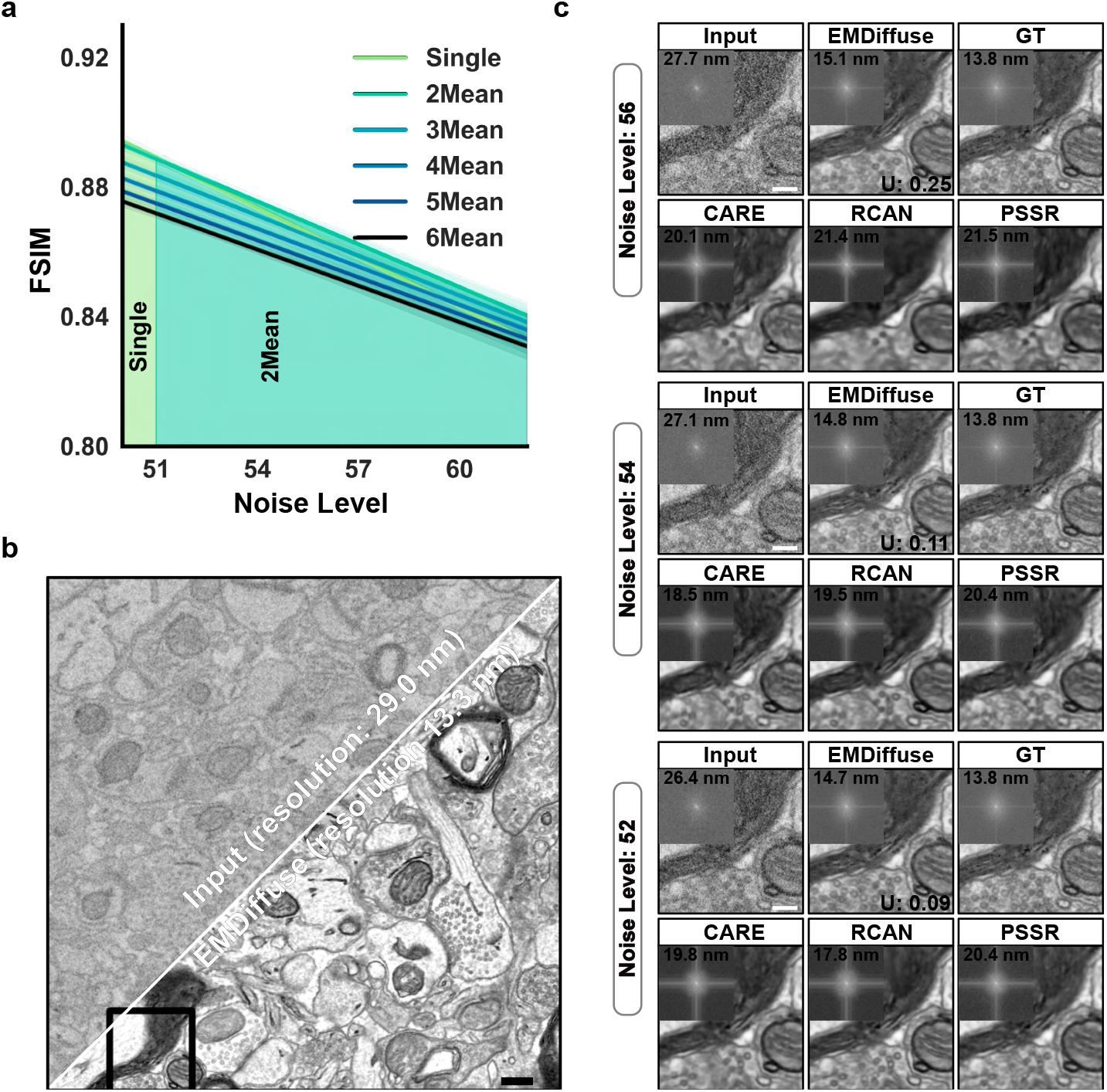
EMDiffuse outperforms other super-resolution methods across various noise levels in structural and perceptual similarities. **a**. FSIM metrics of EMDiffuse super-resolution from different noise levels and different mean numbers on the super-resolution dataset. EMDiffuse adaptively selects the appropriate number of images for dynamic mean based on the noise level of the input image to obtain the highest FSIM score. **b**. Additional example of EMDiffuse super-resolution result and the corresponding input image. **c** Representative images of one enlarged region of **b** with GT, different noise levels input and super-resolution predictions from EMDiffuse, CARE, RCAN, and PSSR. Resolution values and Fourier power spectrum are in each panel’s top left corner. The uncertainty values are indicated in EMDiffuse prediction results. Scale bar, (**b**), 0.3 μm; (**c**), 0.1 μm.

**Extended Data Fig. 10.**
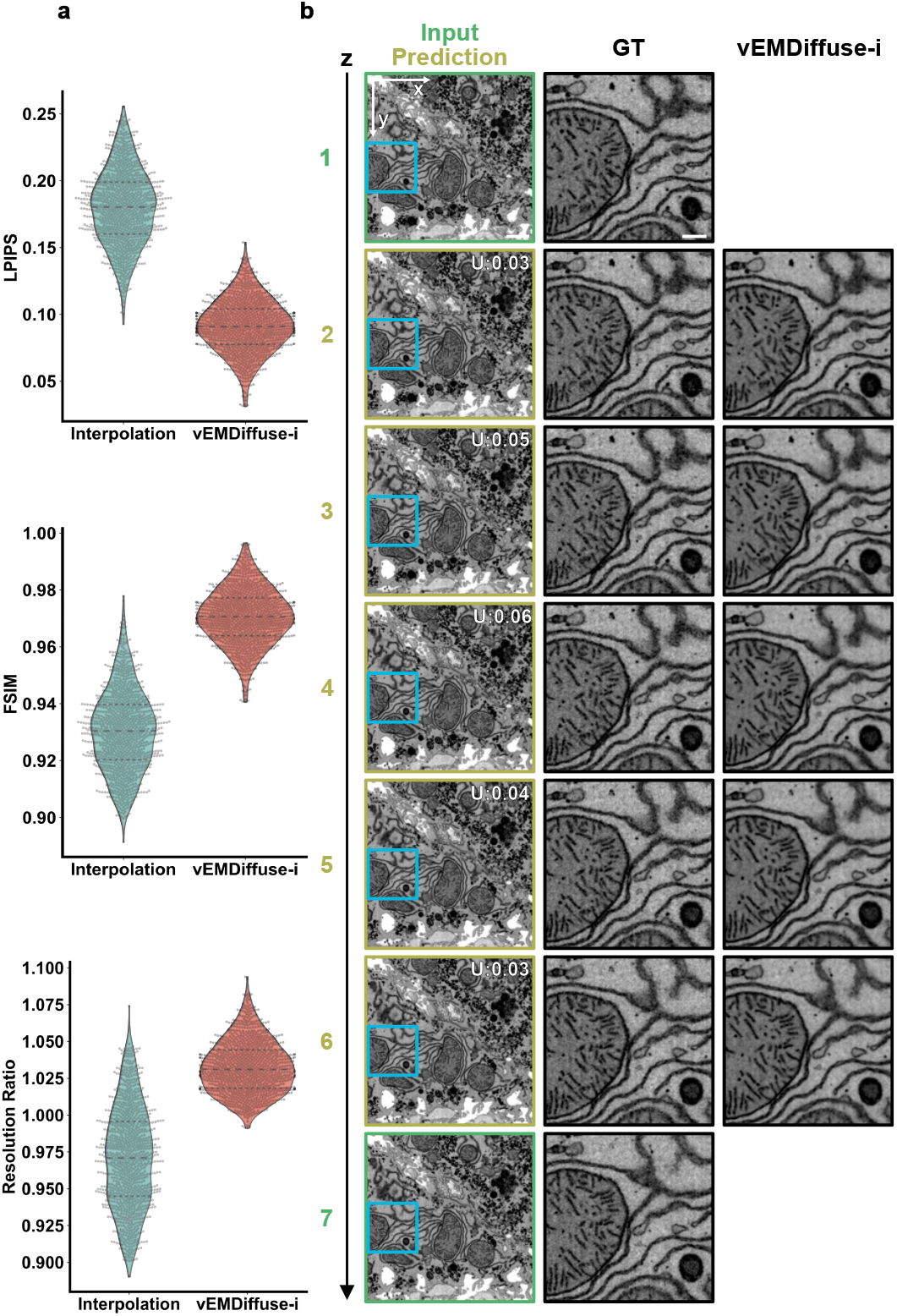
vEMDiffuse-i generates XY Views with isotropic resolution and captures the continuity of organelles along the z-axis in the Openorganelle liver dataset. **a**. The violin plots of three metrics (LPIPS, FSIM, and resolution ratio) show the quantitative performance assessment of vEMDiffuse-i for the isotropic reconstruction task compared with ITK-cubic interpolation. **b**. A series of input and predicted XY views along the z-axis of a liver ultrastructure dataset (jrc_mus-liver). The first column presents the layers of the input (depicted in green) and the vEMDiffuse-i predictions with the uncertainty values (depicted in yellow). The second and third columns showcase layers of the isotropic stack (GT) and vEMDiffuse-i predicted isotropic stack of an enlarged region of the first column. vEMDiffuse-i captures the gradual changes in the ultrastructure of the endoplasmic reticulum (ER), the same as shown in the ground truth slices. Scale bar, first column, 1 μm; second and third columns, 0.3 μm.

**Extended Data Fig. 11.**
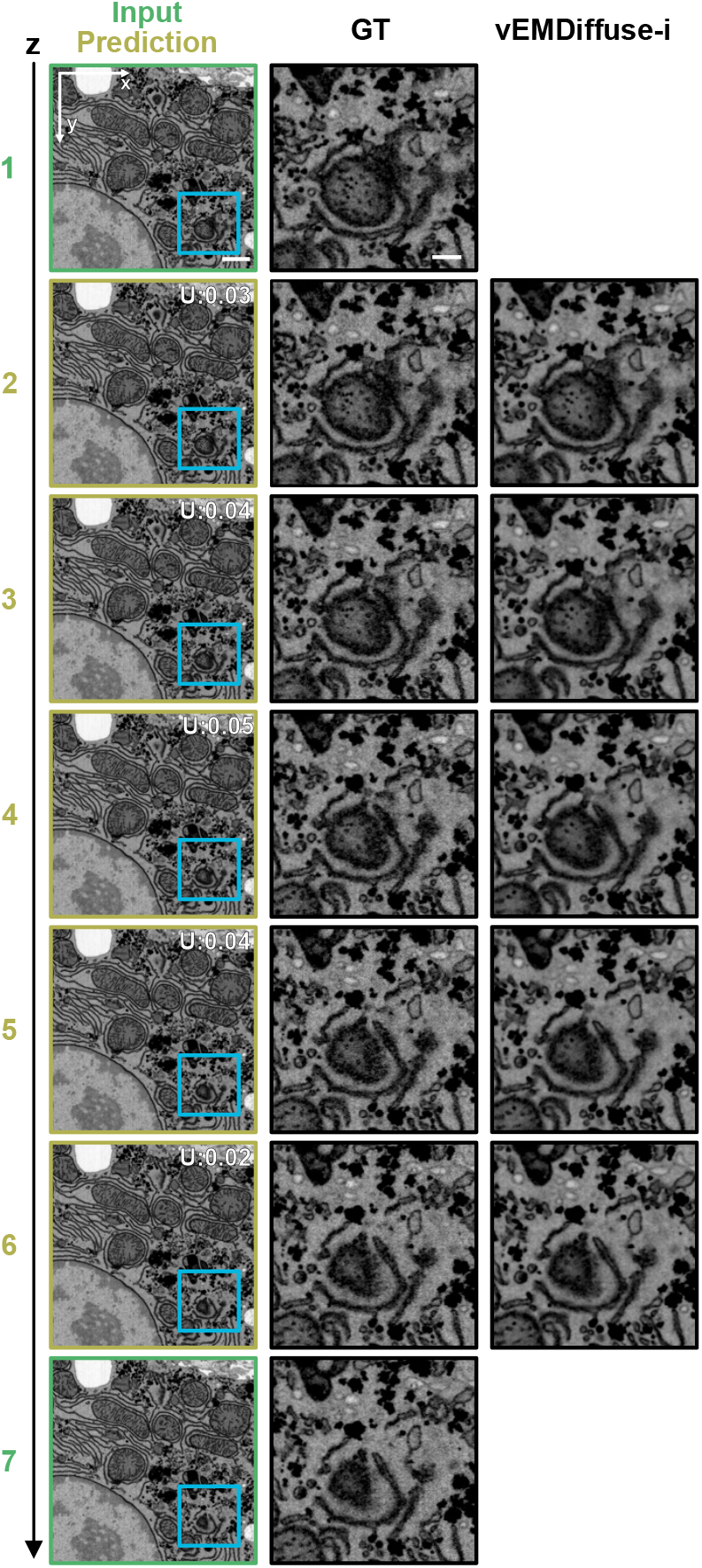
Additional example of the performance of vEMDiffuse-i for generating XY views with isotropic resolution of the Openorganelle liver dataset. Another series of input and vEMDiffuse-i predicted XY views along the z-axis of the Openorganelle liver dataset (jrc_mus-liver). The first column presents the layers of the input (depicted in green) and the vEMDiffuse-i predictions with the uncertainty values (depicted in yellow). The second and third columns showcase layers of the isotropic stack (GT) and vEMDiffuse-i predicted isotropic stack of an enlarged region of the first column. Scale bar, first column, 1 μm, second and third columns, 0.3 μm.

**Extended Data Fig. 12.**
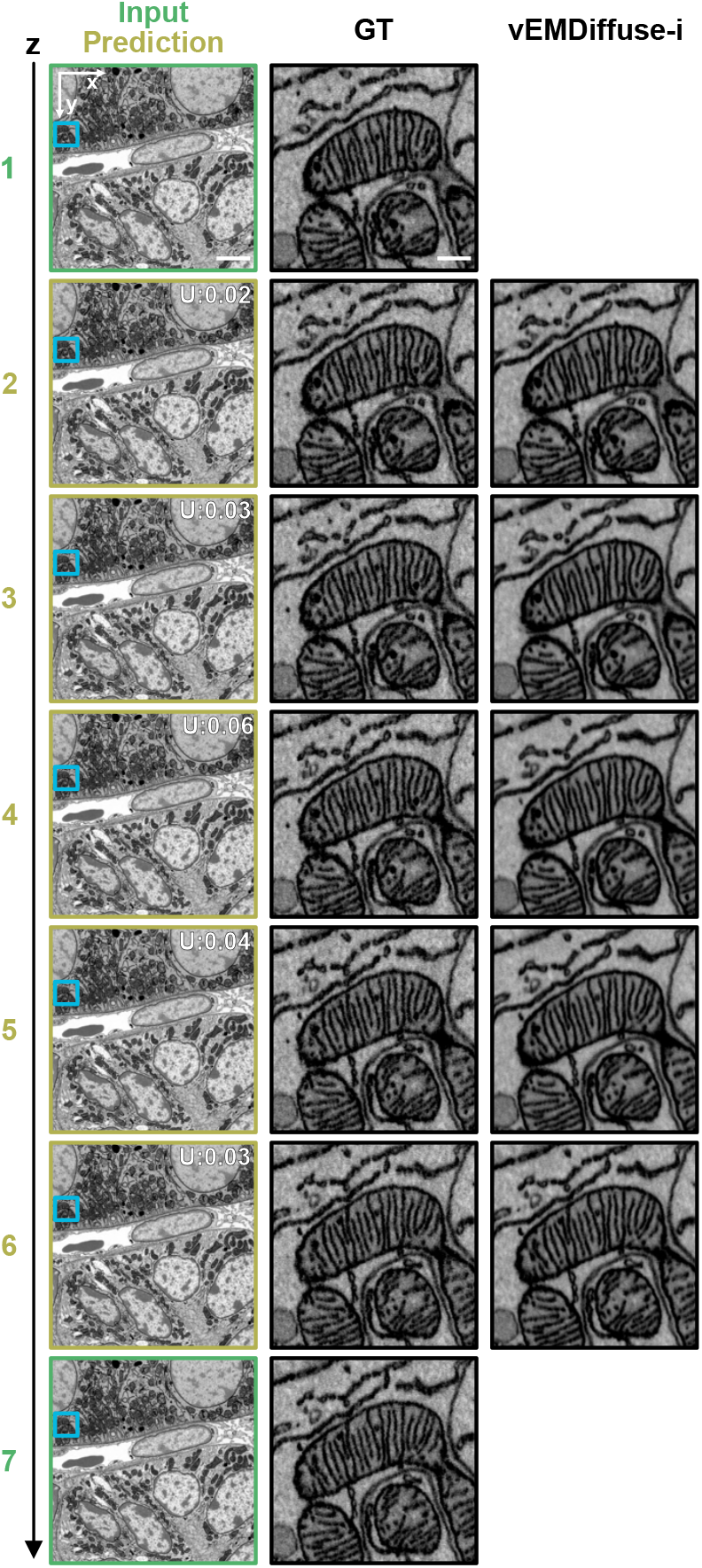
vEMDiffuse-i generates isotropic vEM volumes and captures the continuity of ultrastructure along the z-axis in the Openorganelle kidney dataset. Shown are a series of input and vEMDiffuse-i predicted XY views with uncertainty values along the z-axis with isotropic resolution of the Openorganelle Kidney Dataset (jrc_mus-kidney). The figure is organized identically to Extended Data Fig. 10b and Extended Data Fig. 11. vEMDiffuse-i’s prediction (depicted in yellow) captures the gradual changes of the ultrastructure of the mitochondria and mitochondria cristae as shown in the ground truth. Scale bar, first column, 400 nm; second and third columns, 40 nm.

**Extended Data Fig. 13.**
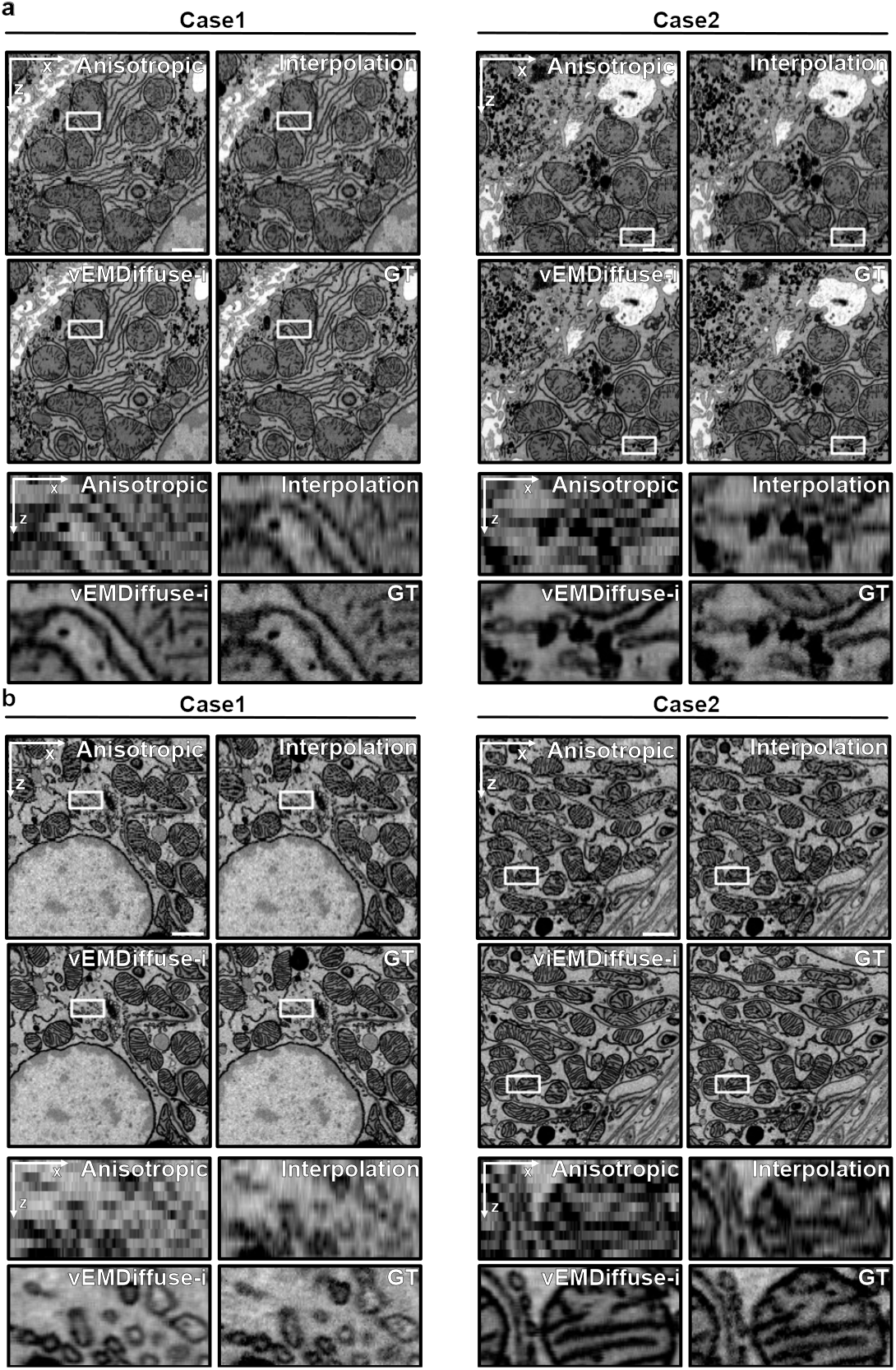
vEMDiffuse-i reconstructs anisotropic volumes into isotropic volumes of two vEM datasets. **a**. XZ views of the ultrastructure of a mouse liver. Shown are two XZ view examples and their enlarged regions of the anisotropic volume, interpolated volume, vEMDiffuse-i reconstructed volume, and isotropic volume (GT) from the Openorganelle liver dataset (jrc_mus-liver). **b**. XZ views of the ultrastructure of a mouse kidney. Shown are two XZ view examples and their enlarged regions of the anisotropic volume, interpolated volume, vEMDiffuse-i reconstructed volume, and isotropic volume (GT) from the Openorganelle kidney dataset (jrc_mus-kidney). Scale bar, 1 μm.

**Extended Data Fig. 14.**
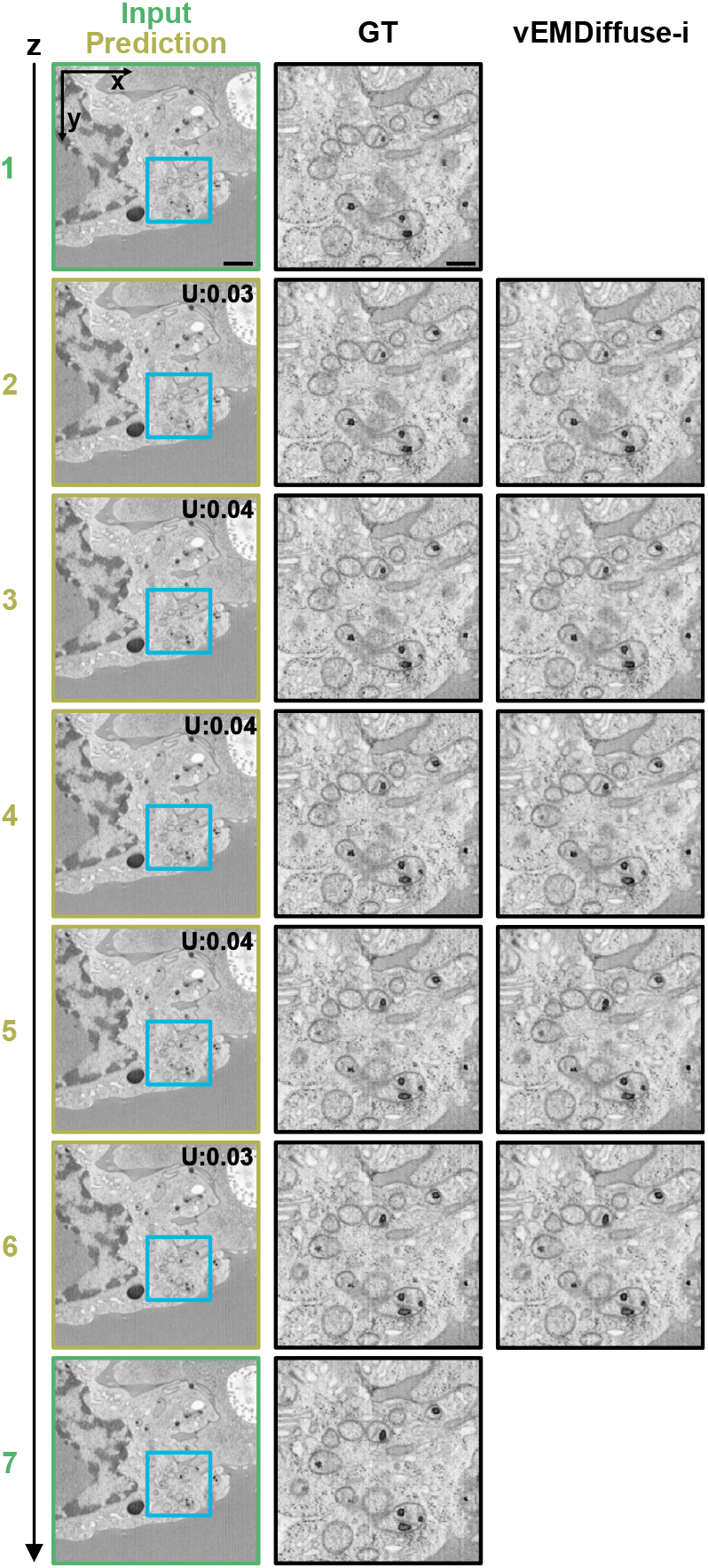
vEMDiffuse-i generates isotropic vEM volumes and captures the continuity of ultrastructure along the z-axis in the T-cell vEM datasets. Shown are a series of input and vEMDiffuse-i predicted XY views with uncertainty values along the z-axis with isotropic resolution of the Openorganelle T-cell Dataset (jrc_ctl-id8-2). The figure is organized identically to Extended Data Fig. 10b, Extended Data Fig. 11 and Extended Data Fig. 12. vEMDiffuse-i’s prediction (depicted in yellow) captures the gradual changes of the ultrastructure of the organelles as shown in the ground truth. Scale bar, first column, 1 μm; second and third columns, 0.3 μm.

**Extended Data Fig. 15.**
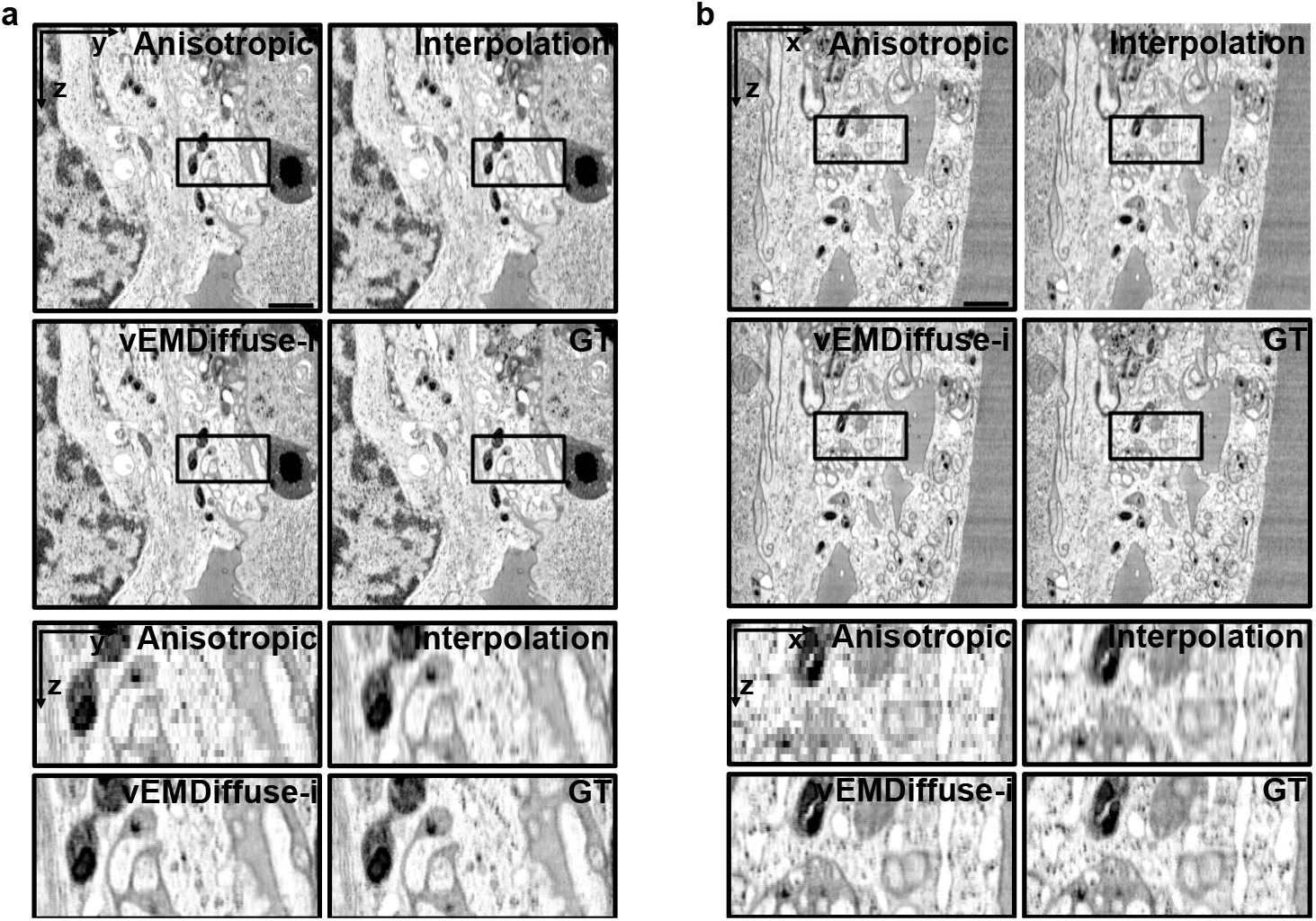
vEMDiffuse-i reconstructs anisotropic volumes into isotropic volumes of T-cell vEM datasets. **a**. YZ view of the ultrastructure of a T-cell. Shown are one YZ view example and one enlarged region of the anisotropic volume, interpolated volume, vEMDiffuse-i reconstructed volume, and isotropic volume (GT) from the Openorganelle T-cell dataset (jrc_ctl-id8-2). **b**. XZ view of the ultrastructure of a T-cell. Shown are one XZ view example and their one regions of the anisotropic volume, interpolated volume, vEMDiffuse-i reconstructed volume, and isotropic volume (GT) from the Openorganelle T-cell dataset (jrc_ctl-id8-2). Scale bar, 1 μm.

**Extended Data Fig. 16.**
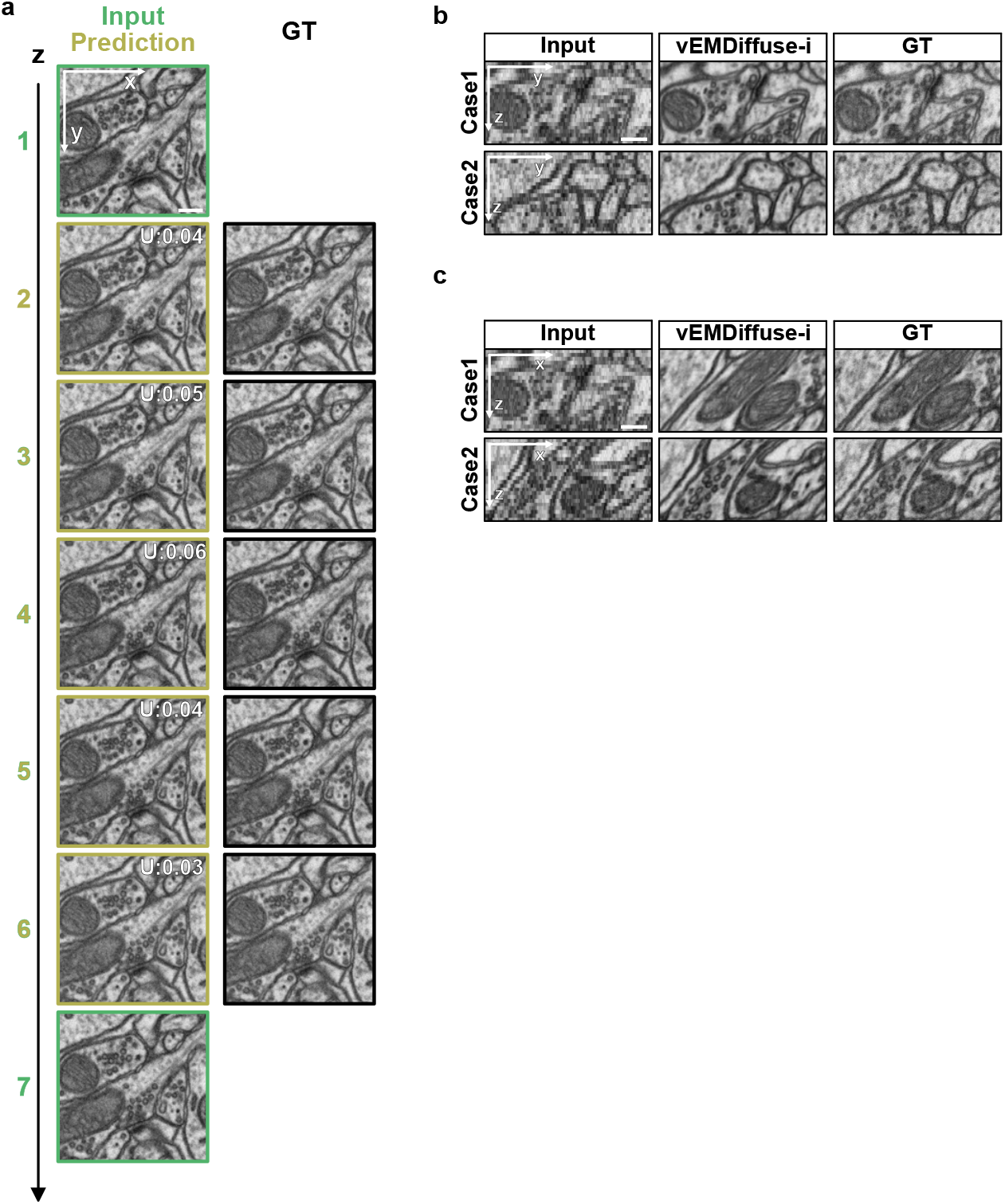
vEMDiffuse-i is transferable for the brain vEM dataset and captures the continuity of neuron ultrastructure. **a**. XY view of vEMDiffuse-i reconstructed brain stack with uncertainty values and isotropic brain stacks from the EPFL brain dataset. The first column presents the layers of the input (depicted in green) and the vEMDiffuse-i predictions with the uncertainty values (depicted in yellow). The second column displays the same layers in ground truth isotropic volume. vEMDiffuse-i captures the gradual appearance of vesicles and discriminates proximal membrane structures in neurons along the z-axis. **b and c**. YZ view (**b**) and XZ view (**c**) of anisotropic brain volume, vEMDiffuse-i generated isotropic volume, and ground truth isotropic volume (GT). Scale bar, (**a**), 0.2 μm; **(b**), 0.1 μm.

**Extended Data Fig. 17.**
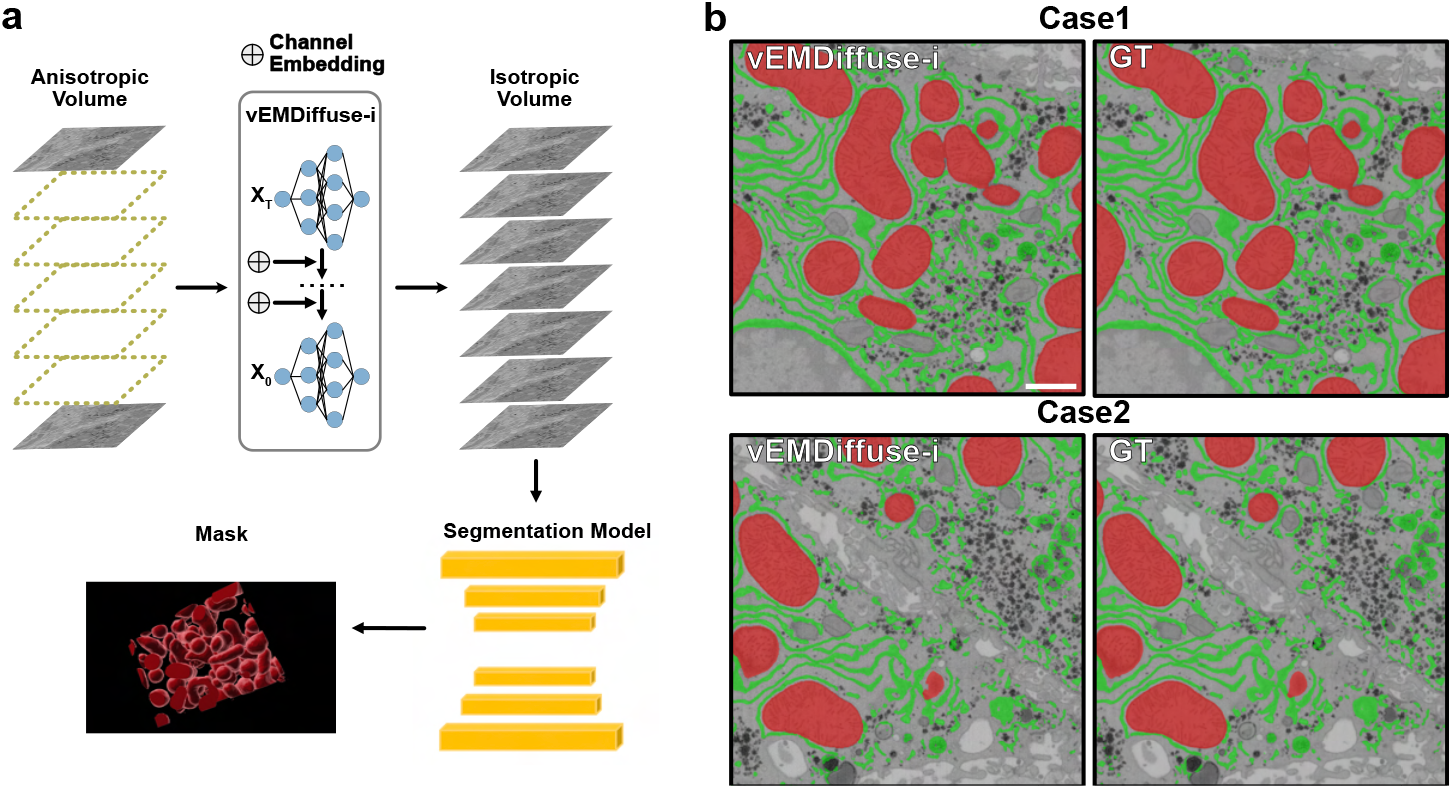
3D reconstruction pipeline and the generation of organelle masks on vEMDiffuse-i reconstructed EM volume and isotropic EM volume. **a**. Schematic of the 3D reconstruction pipeline of isotropic vEM datasets generated by vEMDiffuse-i. A segmentation deep learning model is incorporated to segment organelles of interest for the 3D reconstruction of vEM datasets. **b**. Examples of segmentation results. Two representative XY view composite images of vEMDiffuse-i generated images and corresponding GT images with the organelle segmentation masks generated with the segmentation network. Red mask, mitochondria; green mask, ER. Scale bar, 1 μm.

**Extended Data Fig. 18.**
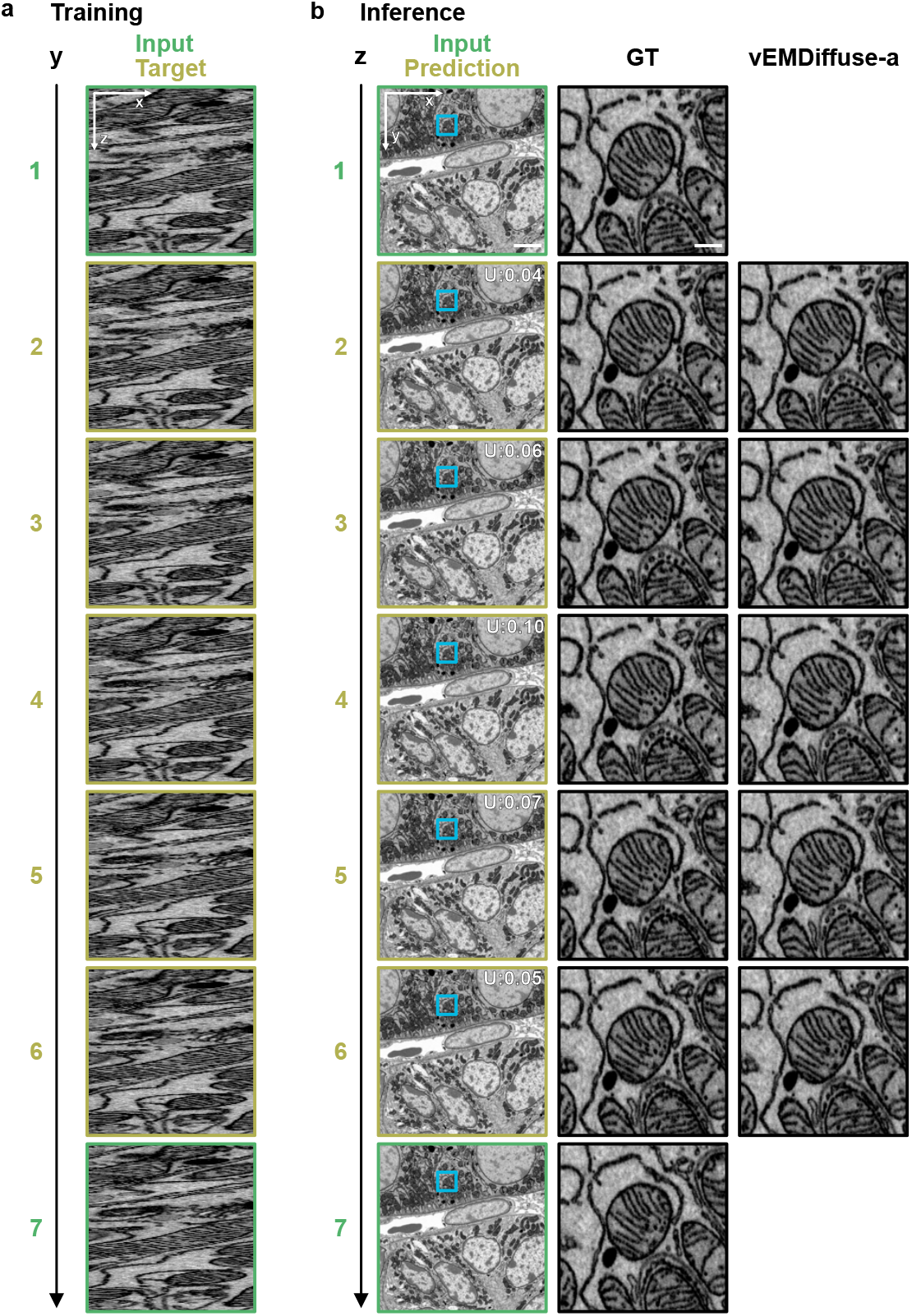
vEMDiffuse-a generates isotropic vEM volumes and captures continuity of ultrastructure along the z-axis with only XZ views as training Data. **a**. Example training images. A series of XZ views with uncertainty values along the y-axis of the downsampled Openorganelle kidney dataset (by removing several z slices) is used to train vEMDiffuse-a. **b**. Example inference images. A series of XY views along the z-axis generated by vEMDiffuse-a.The first column presents the layers of the input (depicted in *green*) and the vEMDiffuse-a predictions (depicted in *yellow*). The second and third columns showcase layers of the isotropic stack (GT) and vEMDiffuse-a predicted isotropic stack of an enlarged region of the first column. vEMDiffuse-a captures the gradual changes in the ultrastructure of the endoplasmic reticulum (ER) and mitochondria, as shown in the ground truth slices. Scale bar, first column, 400 nm; second and third columns, 40 nm.

**Extended Data Fig. 19.**
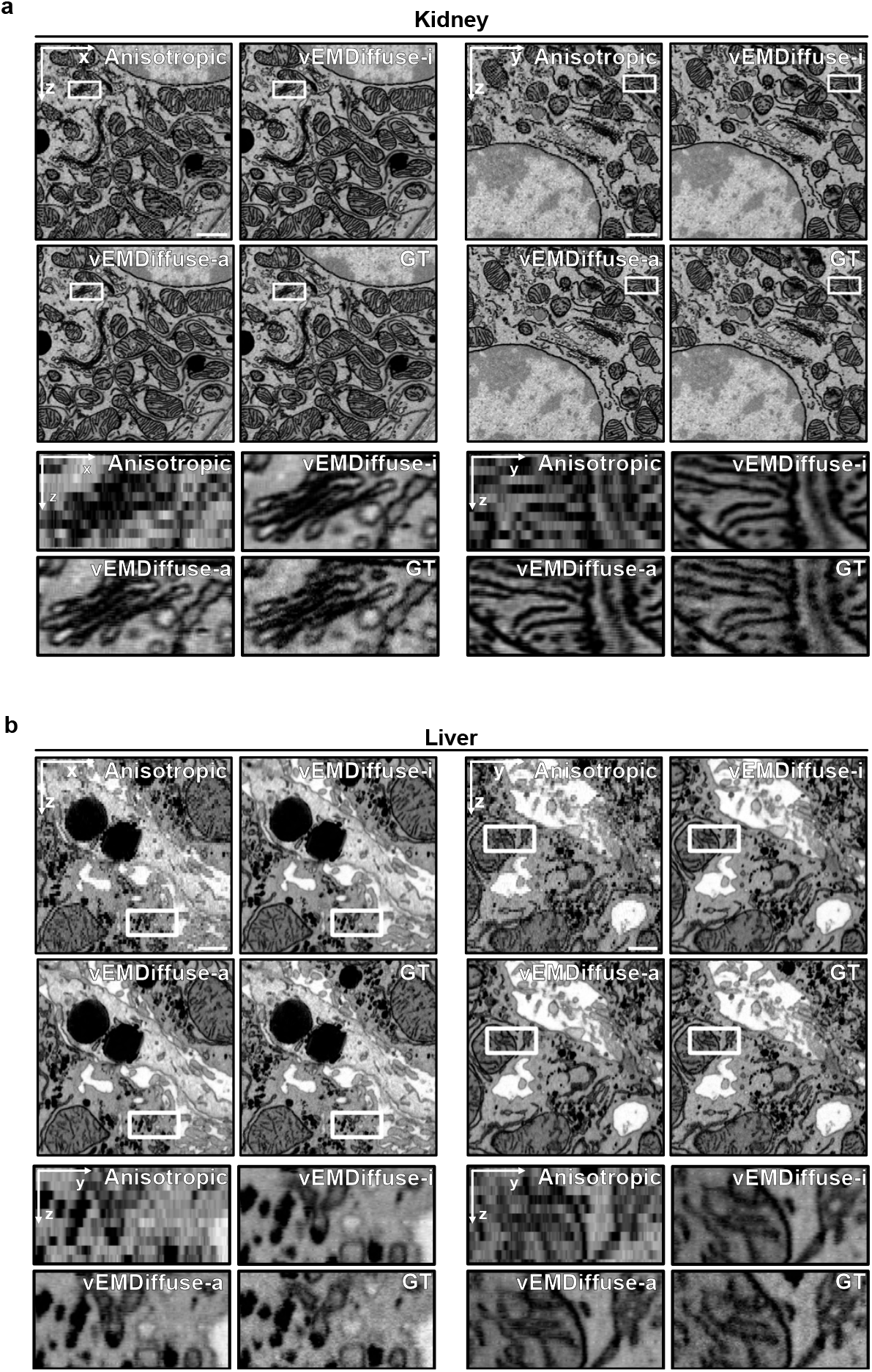
vEMDiffuse-a achieves comparable performance in generating isotropic volumes of vEM datasets with vEMDiffuse-i. Example XZ view and YZ view and enlarged regions of the anisotropic volume, vEMDiffuse-i generated volume, vEMDiffuse-a generated volume, and ground truth isotropic volume (GT) on the Openorganelle kidney dataset (**a**) and Openorganelle liver dataset (**b**). Scale bar, (**a**), 1 μm; **(b**), 0.6 μm.

**Extended Data Fig. 20.**
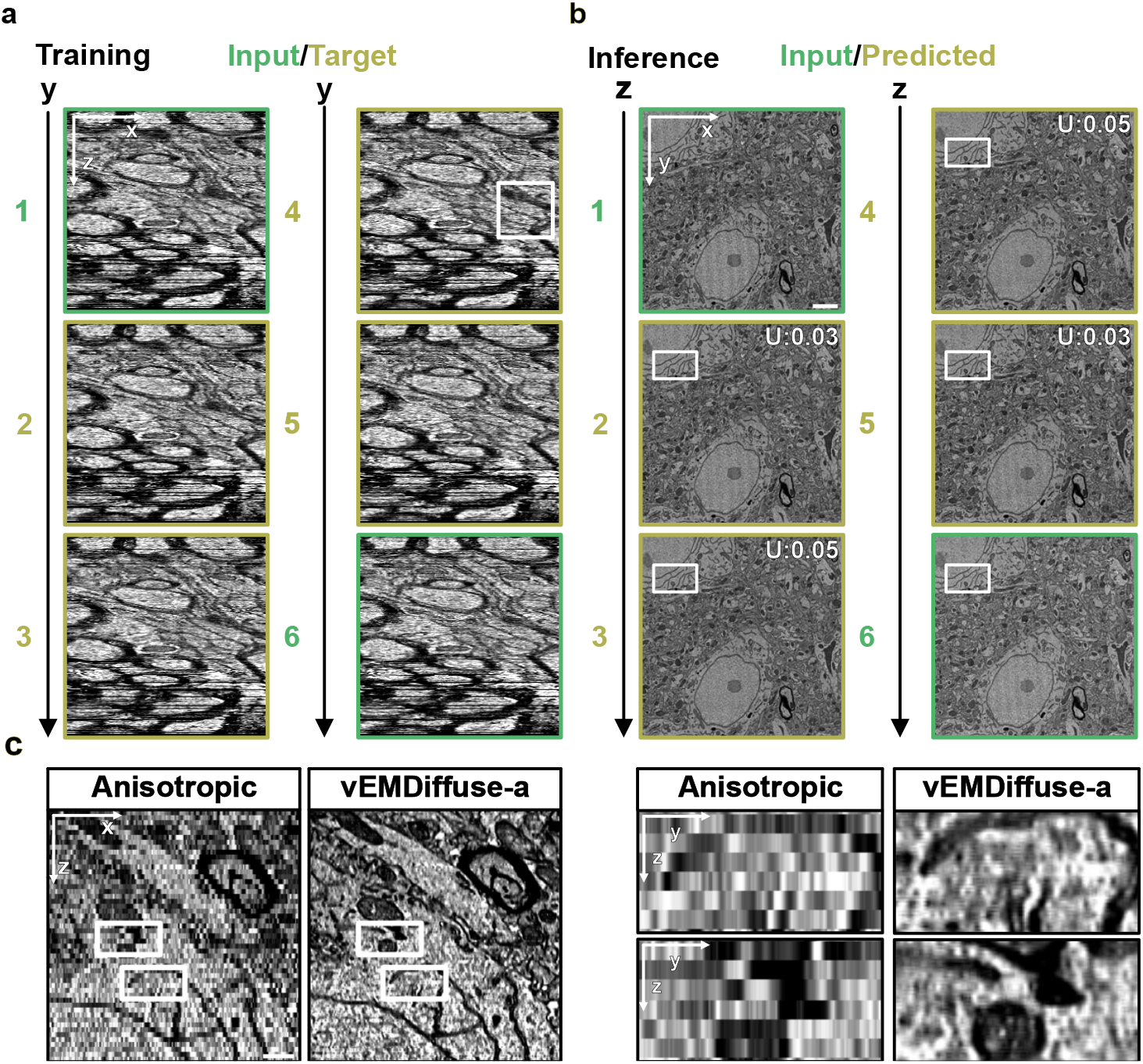
vEMDiffuse-a significantly enhances the axial resolution of the MICrONS multi-area dataset. **a**. Example training images examples. A series of XZ views (input images in *green* and target images in *yellow*) along the y-axis of the anisotropic MICrONS multi-area dataset is used to train vEMDiffuse-a. **b**. Example inference images. A series of vEMDiffuse-a generated XY views (*yellow*) with uncertainty values and input XY views (*green*) along the z-axis. Scale bar, 1 μm. **c**. XZ views and two enlarged regions of the original MICrONS multi-area dataset volume (anisotropic) and vEMDiffuse-a generated volume. Scale bar, (**b**), 2 μm; **(c**), 0.5 μm.

**Extended Data Fig. 21.**
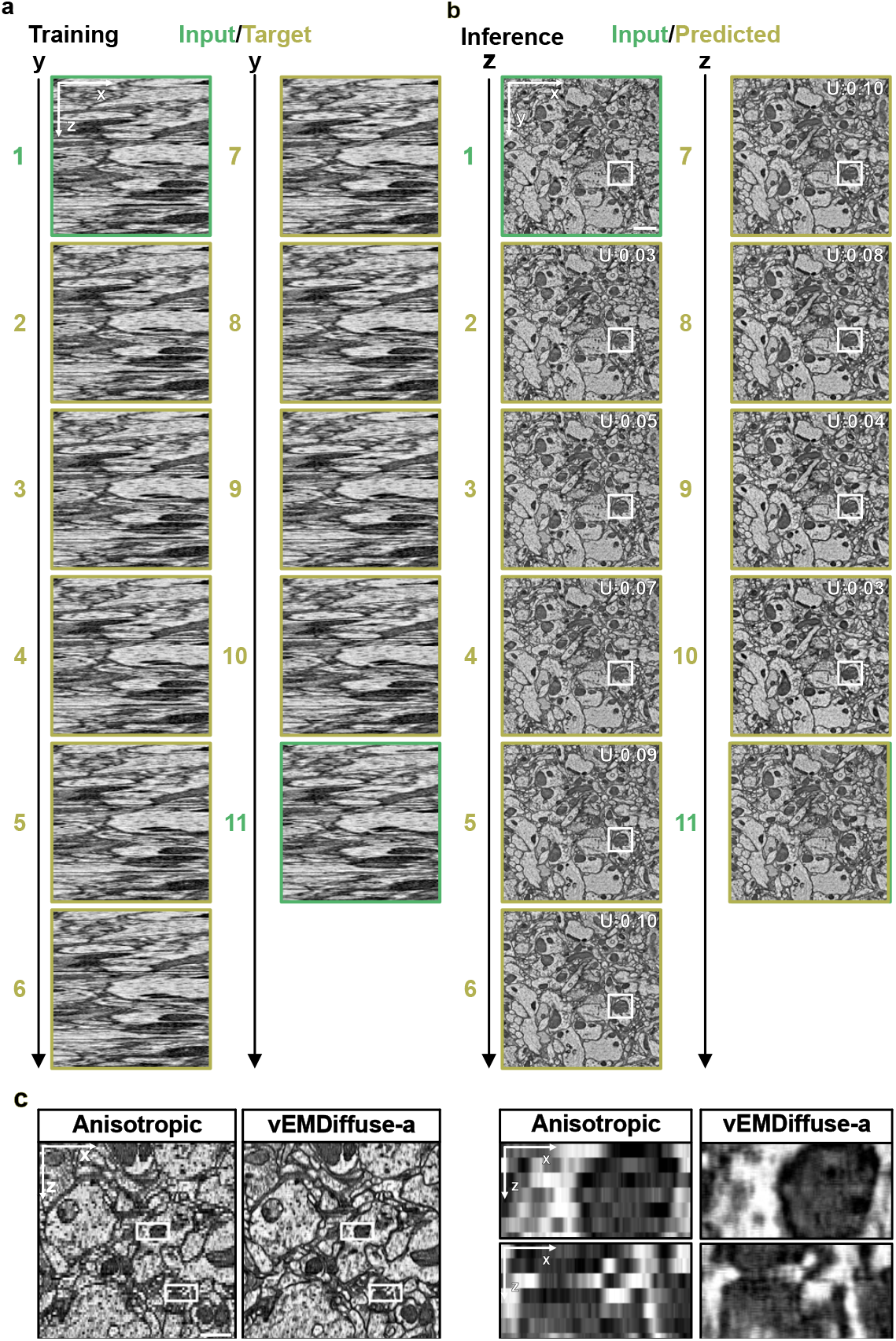
vEMDiffuse-a enhances the axial resolution of the FANC dataset. **a**. Example training images examples. A series of XZ views (input images in *green* and target images in *yellow*) along the y-axis of the anisotropic FANC dataset is used to train vEMDiffuse-a. **b**. Example inference images. A series of vEMDiffuse-a generated XY views (*yellow*) with uncertainty values and input XY views (*green*) along the z-axis. **c**. XZ views and two enlarged regions of the original FANC dataset volume (anisotropic) and vEMDiffuse-a generated volume. Scale bar, (**b**), 1.2 μm; **(c**), 0.5 μm.

## Supplementary Videos

**Supplementary Video 1** Representative illustrative examples of enhanced image quality achieved through EMDiffuse-n denoising on a mouse brain cortex EM image.

**Supplementary Video 2** Comparison of EMDiffuse-n with other denoising methods on an enlarged region of mouse brain cortex EM image. Predictions processed with CARE, PSSR, and RCAN are compared with results obtained with EMDiffuse-n.

**Supplementary Video 3** Transferability of in EMDiffuse denoising. The raw image, image denoised with EMDiffuse-n without fine-tuning, and image denoised with EMDiffuse after fine-tuning are evaluated across datasets from three different tissues: mouse liver, mouse heart, and bone marrow.

**Supplementary Video 4** Representative illustrative examples of super-resolution achieved through EMDiffuse-r on mouse brain cortex EM images.

**Supplementary Video 5** Comparison of EMDiffuse-r with other super-resolution methods on an enlarged region of mouse brain cortex EM image. Results of super-resolution via CARE, PSSR, and RCAN are compared with those achieved through EMDiffuse-r.

**Supplementary Video 6** Examples of transfer learning in EMDiffuse super-resolution. The raw image and image super-resolved with EMDiffuse-r after fine-tuning are evaluated across three distinct tissues: mouse liver, mouse heart, and bone marrow.

**Supplementary Video 7** vEMDiffuse-i restores isotropic volume from downgraded Openorganelle kidney (jrc_mus-kidney) volume. Left: XY views of ultrastructural change within volume reconstructed by vEMDiffuse-i derived from the anisotropic volume with 8 nm x 8 nm x 48 nm voxel size. Right: XY views of ultrastructural change within ground truth isotropic volume with 8 nm x 8 nm x 8 nm voxel size.

**Supplementary Video 8** vEMDiffuse-i restores isotropic volume from downgraded Openorganelle liver (jrc_mus-liver) volume. Left: XY views of ultrastructural change within volume reconstructed by vEMDiffuse-i derived from the anisotropic volume with 8 nm x 8 nm x 48 nm voxel size. Right: XY views of ultrastructural change within ground truth isotropic volume with 8 nm x 8 nm x 8 nm voxel size.

**Supplementary Video 9** 3D reconstruction and rendering of endoplasmic reticulum (ER) and mitochondria within Openorganelle liver volume. Left: ER and mitochondria reconstructions derived from anisotropic volume. Right: ER and mitochondria reconstructions obtained from vEMDiffuse-i restored volume.

**Supplementary Video 10** vEMDiffuse-a restores isotropic volume from downgraded Openorganelle kidney (jrc_mus-kidney) volume without training with isotropic data. Left: XY views of ultrastructural change within volume reconstructed by vEMDiffuse-a derived from the anisotropic volume with 8 nm x 8 nm x 48 nm voxel size. Right: XY views of ultrastructural change within ground truth isotropic volume with 8 nm x 8 nm x 8nm voxel size.

**Supplementary Video 11** vEMDiffuse-a restores isotropic volume from downgraded Openorganelle liver (jrc_mus-liver) volume without training with isotropic data. Left: XY views of ultrastructural change within volume reconstructed by vEMDiffuse-a derived from the anisotropic volume with 8 nm x 8 nm x 48 nm voxel size. Right: XY views of ultrastructural change within ground truth isotropic volume with 8 nm x 8 nm x 8 nm voxel size.

**Supplementary Video 12** Illustrations of YZ views of vEMDiffuse-a enhanced Openorganelle kidney (jrc_mus-kidney) and liver (jrc_mus-liver) volumes.

**Supplementary Video 13** 3D reconstruction and rendering of mitochondria within Openorganelle kidney volume. Yellow: mitochondria reconstructions from anisotropic volume. Red: Organelle reconstruction from vEMDiffuse-a restored volume.

**Supplementary Video 14** vEMDiffuse-a generates isotropic resolution volume from the anisotropic MICrONS multi-area volume. Left: XY views of ultrastructural change within volume generated by vEMDiffuse-a from anisotropic volume. Right: XY views of ultrastructural change in anisotropic volume.

**Supplementary Video 15** vEMDiffuse-a generates isotropic resolution volume from the anisotropic FANC volume. Left: XY views of ultrastructural change within volume generated by vEMDiffuse-a from anisotropic volume. Right: XY views of ultrastructural change in anisotropic volume.

**Supplementary Video 16** Illustrations of YZ views of vEMDiffuse-a generated isotropic FANC and MICrONS multi-area volume.

## Notes

### Competing Interest Statement

The authors have declared no competing interest.

